# Identification of differentially distributed gene expression and distinct sets of cancer-related genes identified by changes in mean and variability

**DOI:** 10.1101/2021.02.15.431343

**Authors:** Aedan G. K. Roberts, Daniel R. Catchpoole, Paul J. Kennedy

## Abstract

There is increasing evidence that changes in the variability or overall distribution of gene expression are important both in normal biology and in diseases, particularly cancer. Genes whose expression differs in variability or distribution without a difference in mean are ignored by traditional differential expression-based analyses. Using a Bayesian hierarchical model that provides tests for both differential variability and differential distribution for bulk RNA-seq data, we report here an investigation into differential variability and distribution in cancer. Analysis of eight paired tumour–normal datasets from The Cancer Genome Atlas confirms that differential variability and distribution are able to identify cancer-related genes. We further demonstrate that differential variability identifies cancer-related genes that are missed by differential expression analysis, and that differential expression and differential variability identify functionally distinct sets of genes. These results suggest that differential variability analysis may provide insights into genetic aspects of cancer that would not be revealed by differential expression, and that differential distribution analysis may allow for more comprehensive identification of cancer-related genes than analyses based on changes in mean or variability alone.

## Background

As RNA sequencing (RNA-seq) has replaced microarray as the leading technology for large-scale gene expression analysis at the whole tissue level, there has been rapid progress in the development of methods for analysing the resulting count data. The focus of most of these methods is differential expression analysis – identifying genes whose mean expression levels differ between groups of interest. However, there is a growing body of evidence to suggest that differences in variability of gene expression are also biologically and medically important. Differences in expression variability have been associated with biological function [1, 2, 3], development [4, 5] and ageing [6, 7, 8, 9]. Changes in expression variability have also been implicated in diseases including schizophrenia [3, 10] and cancer [11, 12, 13, 14]. Genes selected for differences in variability have been demonstrated to have diagnostic and prognostic potential in cancer [12, 13, 15, 16, 14, 17].

Work on expression variability to date has often focused on microarray data, and variability has been assessed using empirical measures such as standard deviation [13, 15, 5], the coefficient of variation (CV; the ratio of the standard deviation to the mean) [11, 3, 10], or measures of deviation from expected expression values determined by a relationship between the mean and variability [18, 17]. Tests for changes in variability have generally been based on established normal distribution-based tests such as the *F*-test and Bartlett’s test, or robust alternatives such as Levene’s test or the Brown–Forsyth test [6, 19, 20, 21, 14, 22]. In assuming that the data follows a normal distribution or not assuming any parametric distribution, these tests are likely to be under-powered for all but very large sample sizes. This is particularly the case if they are to be applied to RNA-seq data, which takes the form of counts, and so is discretely, rather than continuously, distributed.

RNA-seq data is most commonly modelled using the negative binomial (NB) distribution, for which the variance is a function of the mean and a dispersion parameter. There is one method published to date that provides a test for differences in variability specifically for bulk RNA-seq data: MDSeq [23], which uses an NB generalised linear model to test for differences in mean and dispersion separately. Generalised additive models for location, shape and scale (GAMLSS) [24] form another family of regression models with potential for identifying differentially variable genes, as has recently been demonstrated [25]. BASiCS [26] uses a Bayesian model to test for differences in mean and dispersion for single-cell RNA-seq data which relies on spike-in genes or technical replicates to decompose the observed variability into technical and biological components.

The NB distribution is completely defined by the mean and dispersion. “Differential distribution” – any change in the distribution of expression levels between groups – can therefore be defined as a difference in either one or both of these parameters. Since there is a clear interest in identifying genes with differences in mean, and growing evidence for biological relevance of differences in variability, a method that combines both types of analysis into a single test for differential distribution should allow for more complete identification of genes with relevance to a disease or biological state than either method alone.

We have developed a hierarchical Bayesian model based on the NB distribution that provides tests for differential expression, dispersion and distribution for RNA-seq data. The tests for differential expression and dispersion are similar to those implemented in BASiCS. The hierarchical model (HM) is incorporated into a hierarchical mixture model (HMM) which provides an overall test for differential distribution.

Using this model, we report here an investigation into differential variability and distribution in human cancers. Analysis of eight paired tumour–normal RNA-seq datasets from The Cancer Genome Atlas (TCGA; https://www.cancer.gov/tcga) demonstrates that differential variability identifies different sets of genes from differential expression. Using lists of genes previously identified as being related to each of these cancers, we provide a demonstration that differential variability and differential distribution analyses are able to identify cancer-related genes, and that differential variability identifies cancer-related genes that are not identified by differential expression. We further show using gene set enrichment analysis that differential variability identifies functionally distinct sets of genes compared to differential expression.

Together, these results add to the growing body of literature highlighting the importance of changes in expression variability in cancer, and suggest that differential distribution analysis may provide a more comprehensive way of identifying potential cancer-related genes.

## Results

### Hierarchical model identifies genes with differences in mean, dispersion or distribution

We perform differential dispersion and distribution analyses using a Bayesian hierarchical model for RNA-seq data based on the NB distribution. A Markov chain Monte Carlo (MCMC) algorithm provides samples from the posterior distributions of the mean and dispersion for each gene. These posterior samples are the basis for tests for differences in mean or dispersion between groups, and a two-component mixture model provides the basis for a test for differential distribution. The hierarchical nature of the model allows information to be shared among genes, producing parameter estimates for each gene that are shrunk towards a common value estimated over all genes and thereby allowing stable parameter estimation for small sample sizes. Related information-sharing schemes are common in dispersion estimation procedures for differential expression analysis methods [27, 28, 29], and are at the core of Bayesian methods for differential expression analysis [30, 31], but are not used in MDSeq, the only previously published method for differential dispersion in bulk RNA-seq data. Figure 1A shows a schematic outline of the hierarchical model. Further details on the hierarchical model, MCMC sampling and posterior inference are given in Methods.

**Figure 1:**
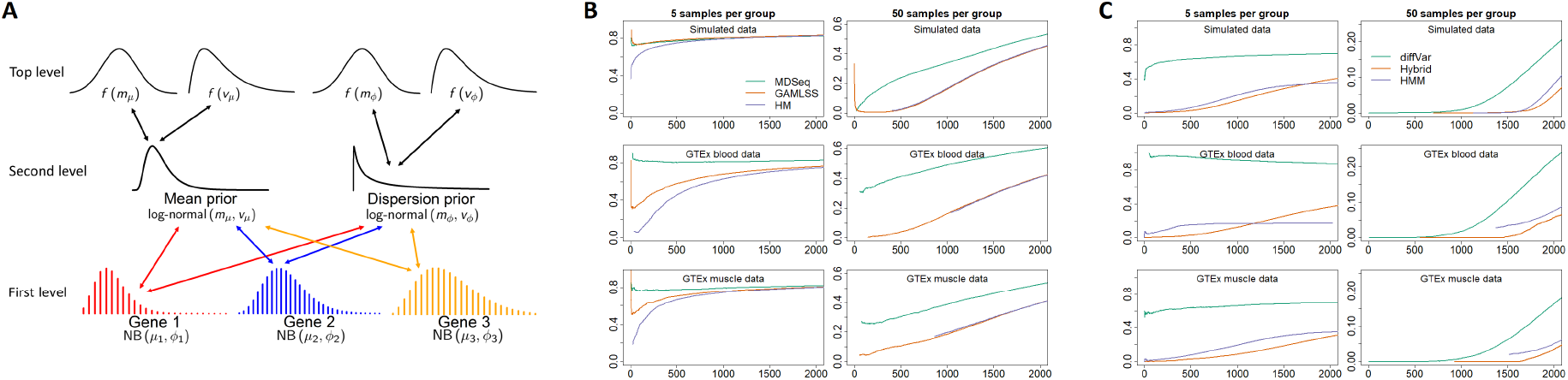
Hierarchical model for differential expression, dispersion and distribution. (A) Schematic illustration of the model. (B) False discovery curves for detection of differential dispersion using the hierarchical model (HM), MDSeq and GAMLSS, averaged over 50 simulated datasets (top) and 10 datasets generated from GTEx blood (middle) and muscle (bottom) data. (C) False discovery curves for differential distribution using the hierarchical mixture model (HMM), diffVar and a hybrid method combining separate tests for differential expression and differential dispersion, averaged over 50 simulated datasets (top) and 10 datasets generated from GTEx blood (middle) and muscle (bottom) data.

Model performance was assessed using two approaches: testing on simulated data, and on real RNA-seq data with artificially-induced differences in expression. Using simulated data allows assessment of the performance of a model with reference to a known ground truth, but has the disadvantage that the data may not adequately reflect the properties of real data. This is an issue especially when the data is simulated under an assumed parametric model – in this case, the NB distribution. An alternative to using simulated data is to use real data, but, with the exception of spike-in experiments, this has the major disadvantage that the ground truth is unknown.

Here we take a different approach: using real data where there is a reasonable assumption of no differences in distribution between groups, and manipulating the data to artificially introduce differences in distribution for a known subset of genes. The Genotype–Tissue Expression (GTEx) project [32] (https://gtexportal.org/home/) provides a source of RNA-seq data from a range of tissues from healthy donors. Randomly splitting samples for a given tissue type into two groups provides a baseline dataset with no expected differences in expression between groups. Using data from two tissues – whole blood and skeletal muscle – we generated datasets with known levels of differential expression and/or dispersion by randomly selecting samples and altering the counts for a proportion of genes in one group to reflect a change in mean and/or dispersion, under the assumption that the counts follow an NB distribution. Full details are given in Methods.

The HM differential expression test was compared against edgeR [33], DESeq2 [34], limma-voom [35, 36], baySeq [30] – which also uses a Bayesian hierarchical model – and MDSeq. HM provided similar performance to the best of the other methods (Additional file 1: Section S1).

The HM differential dispersion test was compared against MDSeq and GAMLSS. False discovery curves (Figure 1B) show that HM consistently outperforms MDSeq, and outperforms GAMLSS with 5 samples per group while providing similar performance with 50 samples per group. False discovery curves for 2, 10 and 20 samples per group (Additional file 1: Figure S3) show similar patterns, as do boxplots of false discovery rates (FDR) and sensitivity (Additional file 1: Figure S4).

Along with MDSeq, HM has an advantage over GAMLSS in being able to test for differences in dispersion at a given minimum log fold change (LFC). False discovery curves (Additional file 1: Figure S5) and boxplots of FDR and sensitivity (Additional file 1: Figure S6) show that HM clearly outperforms MDSeq for differential dispersion at a minimum LFC of 1.44.

Along with the HMM differential distribution test, an alternative method of testing for differential distribution was also considered: a “hybrid” method combining separate tests for differential expression and differential dispersion (see Methods). While there are no published methods for differential distribution for RNA-seq data, another alternative was considered taking advantage of the fact that under the NB model, the variance is function of the mean and the dispersion. This means that a test for a difference in variance should effectively act as a test of differential distribution. As such, diffVar [37, 38] was also included as an alternative test of differential distribution.

False discovery curves (Figure 1C) show that both HMM and hybrid tests are able to detect differentially distributed genes, and that both are more effective than diffVar, particularly for small sample sizes. With 50 samples per group, HMM and the hybrid method identify up to around 1900 differentially distributed genes while maintaining the FDR below 0.05. False discovery curves for 2, 10 and 20 samples per group are shown in Additional file 1: Figure S7, and boxplots of FDR and sensitivity in Additional file 1: Figure S8.

### Differential variability and differential distribution identify cancer-related genes

To test whether differential variability and distribution analyses are able to identify cancer-related genes, we applied tests of differential expression, dispersion and distribution to paired normal-tumour RNA-seq data from eight cancer types from TCGA, and compiled lists of genes that have previously been identified as being related to each cancer from five different databases: the Cancer Genome Census (CGC) [39], DisGeNET [40], IntOGen [41], the Kyoto Encyclopedia of Genes and Genomes (KEGG) [42] and Malacards [43]. We then tested the ability of each method to rank cancer-related genes above other genes using Wilcoxon rank-sum tests. The methods included were differential expression (using limma–voom), differential dispersion using HM, and differential distribution using both HMM and the hybrid method combining limma–voom for differential expression and the HM differential dispersion test.

The resulting *p*-values are given in Table 1. Cancer-related genes are statistically significantly ranked above other genes at the 0.05 level for all eight cancers by differential expression, and for three out of eight by differential dispersion: thyroid carcinoma, lung adenocarcinoma and lung squamous cell carcinoma. For differential distribution, statistically significant associations were identified for all cancers using the hybrid method, and for lung adenocarcinoma, hepatocellular carcinoma and lung squamous cell carcinoma using HMM. This difference can be explained by the extremely small *p*-values that limma–voom returns for many differential expression tests: around half of all *p*-values returned by limma–voom are smaller than the minimum tail probability from the HM differential dispersion test, meaning that the hybrid method is effectively biased towards differentially expressed genes.

**Table 1:** TCGA tumour–normal sample pairs and *p*-values from Wilcoxon rank-sum tests for ranking cancer-related genes above other genes

| TCGA project ID and ref | TCGA project name | Sample pairs | Differential expression | Differential dispersion | Differential distribution (HMM) | Differential distribution (Hybrid) |
| --- | --- | --- | --- | --- | --- | --- |
| BRCA [44] | Breast invasive adenocarcinoma | 182 | $9 \times 10^{-13}$ | 1.00 | 0.16 | $5 \times 10^{-11}$ |
| KIRC [45] | Kidney renal clear cell carcinoma | 144 | $5 \times 10^{-4}$ | 0.94 | 0.08 | $3 \times 10^{-4}$ |
| THCA [46] | Thyroid carcinoma | 98 | 0.003 | 0.002 | 0.28 | $9 \times 10^{-4}$ |
| LUAD [47] | Lung adenocarcinoma | 86 | 0.009 | $1 \times 10^{-4}$ | 0.005 | $1 \times 10^{-4}$ |
| LIHC [48] | Liver hepatocellular carcinoma | 98 | $1 \times 10^{-12}$ | 0.52 | 0.005 | $4 \times 10^{-10}$ |
| LUSC [49] | Lung squamous cell carcinoma | 92 | $3 \times 10^{-4}$ | $3 \times 10^{-4}$ | 0.003 | $3 \times 10^{-5}$ |
| PRAD [50] | Prostate adenocarcinoma | 92 | $1 \times 10^{-10}$ | 0.17 | 0.91 | $1 \times 10^{-10}$ |
| COAD [51] | Colon adenocarcinoma | 78 | $2 \times 10^{-17}$ | 1.00 | 0.89 | $4 \times 10^{-11}$ |

While differential expression analysis ranks previously identified cancer-related genes above other genes more strongly than differential dispersion or distribution in most cases, this should not be surprising, since differential expression has been one of the main methods used to identify cancer-related genes. Given this, it is particularly striking that for three of the eight cancer types, there are similar levels of evidence for association with cancer-related genes for differential dispersion and differential expression.

### Differential variability identifies different sets of genes from differential expression

We next asked whether differential variability and differential expression identify different sets of genes. This is a critical question, and one that has not been addressed comprehensively in previous studies: if differential variability identifies the same sets of genes as differential expression, the method is of little benefit.

To address this, we assessed the correlation between gene lists identified by differential expression and differential dispersion tests. For a given *p*-value or posterior probability threshold for calling a gene as differentially expressed or dispersed, we created a ranked list of all genes called by both methods and calculated the Spearman correlation – the correlation of the ranks – between the two methods. For a given threshold, a positive correlation means that differential expression and differential dispersion are producing similarly ranked lists of genes, while zero or negative correlation means that the two methods are producing unrelated or inversely associated gene rankings.

Results are shown in Figure 2A for lung and prostate adenocarcinoma, and in Additional file 1: Figure S9 for all eight cancers. Negative correlations are observed for most of the range of thresholds, and correlations are more strongly negative for the most highly ranked genes. Additional file 1: Figure S9 also shows corresponding *p*-values from hypothesis tests for negative correlation, with the null hypothesis of zero or positive correlation rejected for all thresholds that are likely to be of practical interest for all eight cancers. These results clearly show that differential expression and differential dispersion identify distinct sets of genes.

**Figure 2:**
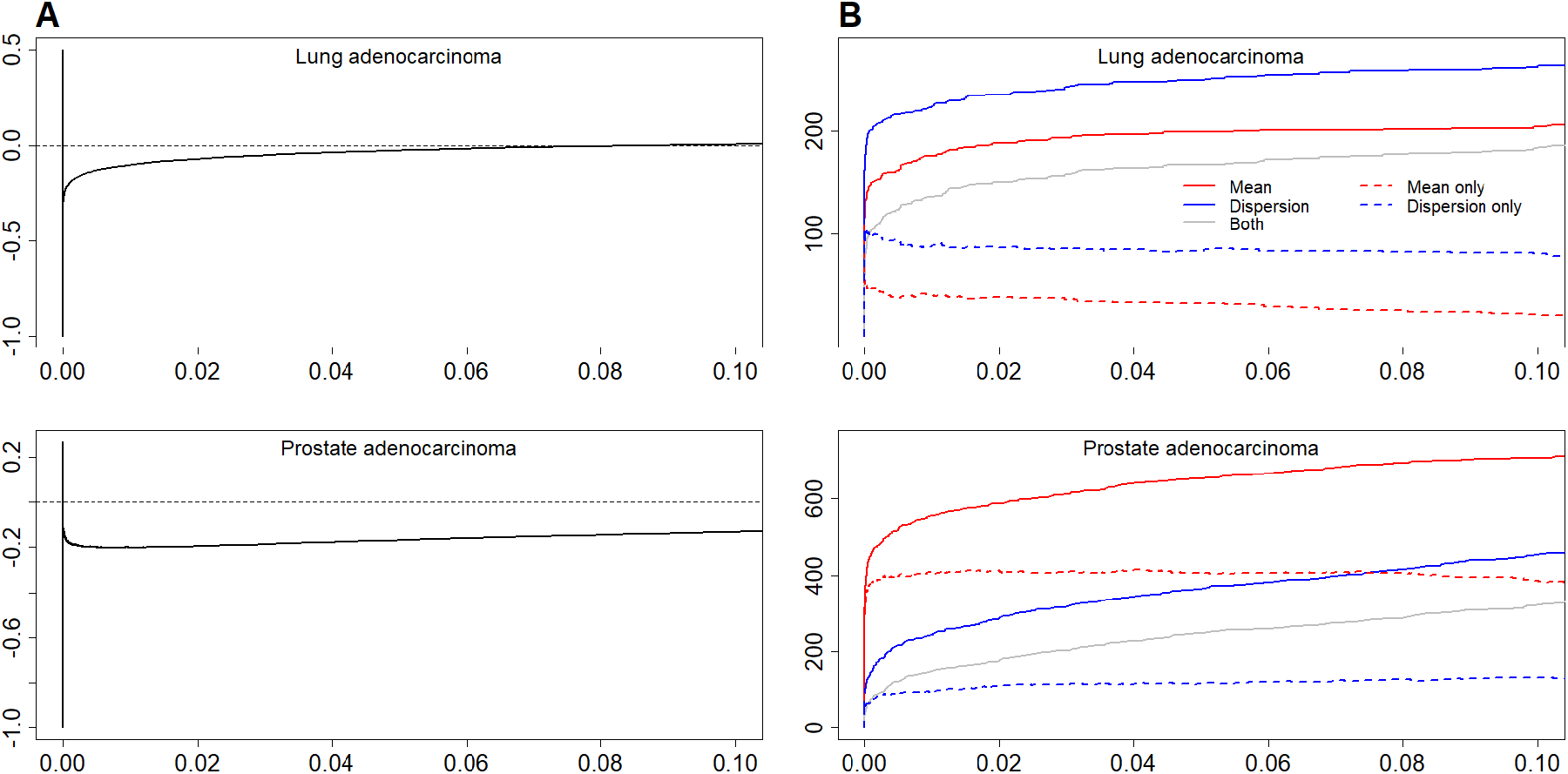
Differential expression and differential dispersion analyses independently identify cancer-related genes. (A) Spearman correlation between gene lists for differential expression and differential dispersion for TCGA lung and prostate adenocarcinoma data, with varying threshold for calling differential expression or dispersion. (B) Number of lung and prostate adenocarcinoma-related genes identified with varying thresholds for calling differential expression or dispersion. Solid lines show the total number of genes identified by differential expression (red), by differential dispersion (blue), and those identified by both methods (grey). Dashed lines show the number of genes uniquely identified by differential expression (red) and by differential dispersion (blue).

These results and those in the previous section demonstrate that, at least for some cancer types, differential variability analysis is able to identify cancer-related genes and that it identifies different sets of genes from differential expression. We next looked specifically at whether differential expression and differential variability identify different sets of cancer-related genes.

Figure 2B shows, for lung and prostate adenocarcinomas, the number of cancer-related genes identified by each method alone and combined, as the threshold for calling a gene as differentially expressed or dispersed is varied. Results for all eight cancers are given in Additional file 1: Figure S9. The results are consistent with the Wilcoxon rank-sum test results: there are more lung adenocarcinoma-related genes identified by differential dispersion than by differential expression, and the opposite for prostate adenocarcinoma. Notably, however, in both cases, even with high thresholds, there are some cancer-related genes that are identified uniquely by each method. For example, even though there is no overall evidence for an association between gene ranking by differential dispersion and prostate adenocarcinoma-related genes, there are around 100 genes that are correctly identified by differential dispersion but not by differential expression. These results provide further support for the idea that neither differential expression nor differential variability should be relied on alone to identify cancer-related genes, and that overall assessment of differential distribution can more comprehensively identify these genes.

### Differential expression and differential variability identify genes in different functional categories

To further investigate the types of genes identified by differential expression and differential variability, we performed gene set enrichment analysis. We looked for enriched terms in each of the three Gene Ontology (GO) ontologies – biological process, molecular function and cellular component – among genes ranked by differential expression and differential dispersion.

Tables 2 and 3 show the ten most significantly enriched terms in each of the three ontologies for genes ranked by differential expression and differential dispersion, respectively, for lung adenocarcinoma. Some general themes are evident, most notably the high proportion of terms related to the immune system and signalling for differential expression, and to transcription, translation and intracellular transport for differential dispersion. Results for the other cancer types are given in Additional file 1: Section S4 and Additional file 2.

**Table 2:** Top 10 enriched GO terms in each ontology for lung adenocarcinoma based on differential expression

| GO term | GO ID | FDR $q$-value |
| --- | --- | --- |
| *Biological process* |  |  |
| Phagocytosis recognition | GO:0006910 | 0.0000 |
| B-cell mediated immunity | GO:0019724 | 0.0000 |
| Lymphocyte-mediated immunity | GO:0002449 | 0.0000 |
| Defense response to bacterium | GO:0042742 | 0.0000 |
| Phagocytosis | GO:0006909 | 0.0000 |
| Immunoglobulin production | GO:0002377 | 0.0000 |
| Membrane invagination | GO:0010324 | 0.0000 |
| FC receptor-mediated stimulatory signaling pathway | GO:0002431 | 0.0000 |
| Urogenital system development | GO:0001655 | 0.0000 |
| Regulation of body fluid levels | GO:0050878 | 0.0000 |
| *Molecular function* |  |  |
| Antigen binding | GO:0003823 | 0.0000 |
| Immunoglobulin receptor binding | GO:0034987 | 0.0000 |
| Receptor regulator activity | GO:0030545 | 0.0003 |
| Extracellular matrix structural constituent | GO:0005201 | 0.0003 |
| Oxygen binding | GO:0019825 | 0.0003 |
| G-protein coupled receptor activity | GO:0004930 | 0.0004 |
| Tetrapyrrole binding | GO:0046906 | 0.0005 |
| Iron ion binding | GO:0005506 | 0.0006 |
| Carbohydrate binding | GO:0030246 | 0.0006 |
| Glycosaminoglycan binding | GO:0005539 | 0.0006 |
| *Cellular component* |  |  |
| Immunoglobulin complex circulating | GO:0042571 | 0.0000 |
| Extracellular matrix | GO:0031012 | 0.0000 |
| External side of plasma membrane | GO:0009897 | 0.0000 |
| Collagen trimer | GO:0005581 | 0.0000 |
| DNA packaging complex | GO:0044815 | 0.0000 |
| Apical plasma membrane | GO:0016324 | 0.0000 |
| Blood microparticle | GO:0072562 | 0.0000 |
| Anchored component of membrane | GO:0031225 | 0.0001 |
| I band | GO:0031674 | 0.0001 |
| Basolateral plasma membrane | GO:0016323 | 0.0002 |

**Table 3:**
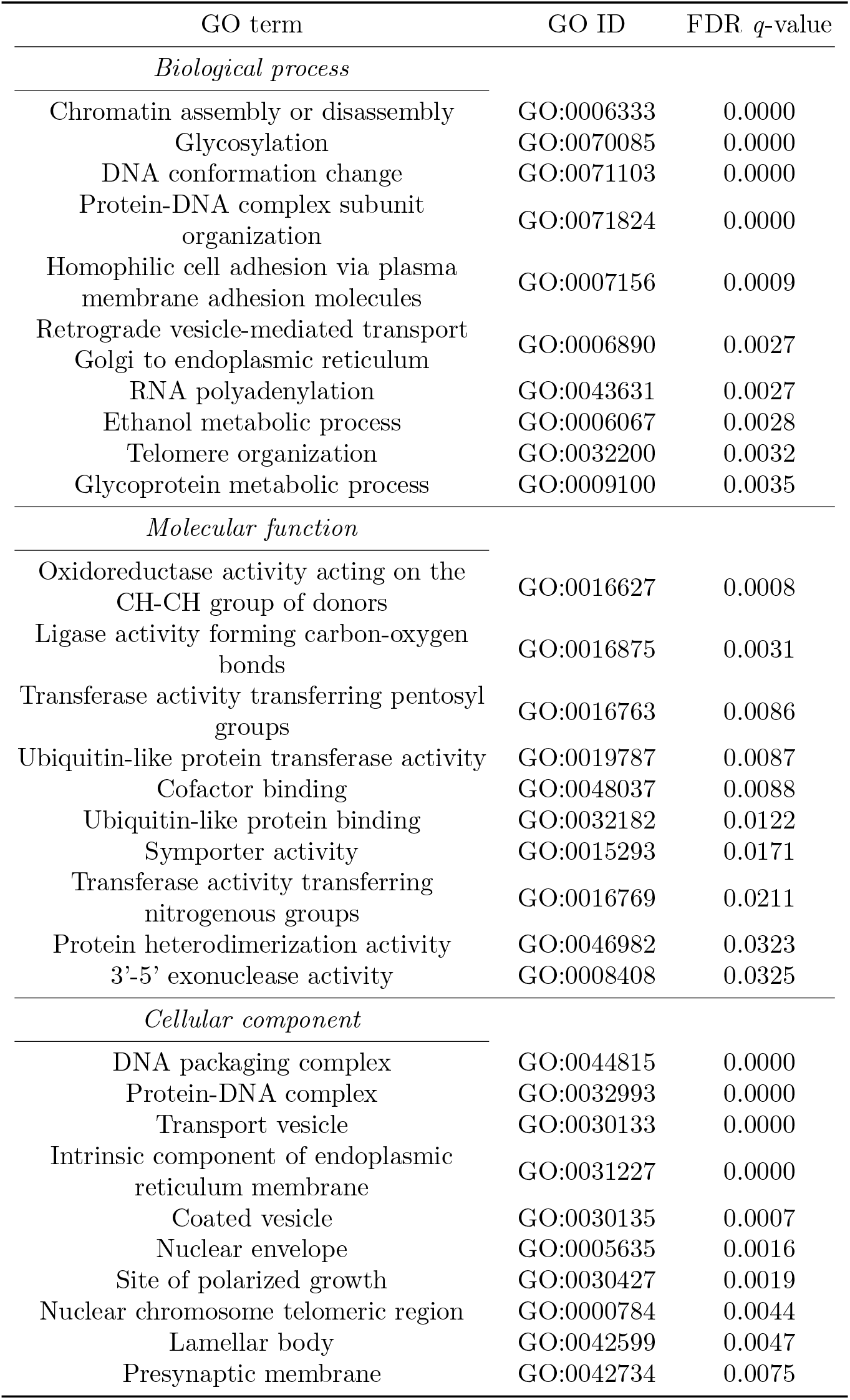
Top 10 enriched GO terms in each ontology for lung adenocarcinoma based on differential dispersion

GO categories relating to DNA replication, transcription and translation are among the most significantly enriched terms for several other cancer types with genes ranked by differential dispersion. For example, for breast adenocarcinoma, RNA 3’-end processing, gene silencing, DNA recombination, telomere organisation and mRNA processing are among the top 10 terms for biological process; histone binding, single-stranded DNA binding, chromatin binding and basal transcription machinery binding are among the top 10 terms for molecular function; and nuclear chromosome telomeric region and replisome are among the five terms with FDR-adjusted q-values < 0.05 for cellular component. In contrast, there are very few such terms among the most significantly enriched with genes ranked by differential expression. To assess the consistency of the GO terms identified using differential variability analysis, we repeated this analysis using GAMLSS instead of the hierarchical model. These results are given in Additional file 2, and show very high concordance between the two differential variability methods.

## Discussion

### Differential variability and distribution in cancer

While there is strong evidence in the existing literature that differentially variable genes are important in cancer, this work provides the first clear demonstration of the inverse: that differential variability can be used to identify cancer-related genes. Importantly, these results also show that differential expression and differential variability identify distinct sets of cancer-related genes. Differential variability analysis ranked previously identified cancer-related genes higher than other genes for three out of eight cancer types tested, with strength of evidence for association similar to that found for differential expression. This is particularly remarkable since it can reasonably be assumed that most cancer-related genes identified to date have been identified at least in part because they have been found to differ in mean expression levels between normal and tumour tissues.

As well as identifying different genes, gene set enrichment analysis showed that differential expression and differential variability identify functionally distinct sets of genes. GO terms relating to basic cellular processes such as transcription and translation are frequently enriched among genes with differences in dispersion between normal and tumour samples. For example, ncRNA metabolic process, nucleic acid phosphodiester bond hydrolysis, RNA 3’-end processing and mRNA processing are all significantly enriched among differentially dispersed genes for multiple cancer types. In contrast, GO terms commonly enriched among differential expressed genes are often related to cell structure and migration (for example extracellular structure organisation), signalling (for example G protein-coupled receptor activity) or immune system functions (for example antigen binding).

Functions relating to transcription and translation have previously been identified among low-variability genes [52, 5, 53]. These are processes that need to be tightly regulated for cells to function properly, and so any change in expression – up or down – for genes involved in these processes is likely to disrupt normal cell function. This suggests a possible biological basis for the differences in types of genes identified by differential expression and differential variability in cancer. Genes identified by differential expression may be either genes that are normally active and for which loss of expression disrupts normal signalling or cellular transport processes, or genes that are normally expressed only in certain cell types or at certain times and for which constitutive expression similarly disrupts normal cell function. On the other hand, genes identified by differential variability may be either genes that under normal circumstances are consistently expressed within a narrow range of levels, the regulation of which is lost in cancer, or genes whose normal function requires changes in expression levels in response to signals, and for which tightening of expression levels therefore disrupts their normal function. This idea is also consistent with previous observations that low variance genes often have “housekeeping” functions, while genes with high expression variability often have functions related to development and response to extracellular signals, for which changes in expression in response to signals are crucial [18]. Altered biological states in cancer may be dependent on different sets of genes being more or less tightly regulated than in healthy tissue, which may result in differences in variability between normal and tumour tissues. Decreased variability among a set of genes associated with invasive potential has previously been observed across multiple cancers [11], with the suggestion that tumour progression may be dependent on the precise regulation of these genes.

There is a distinction between cell-to-cell variability in gene expression at the tissue level and individual-to-individual variability at the population level, the former measured using single-cell RNA-seq and the latter using bulk RNA-seq or microarray. This distinction has not always been made explicit in studies on expression variability, and the arguments above are stronger in the context of differences in expression between cells within a tissue. There have been suggestions that there is some correlation between the different levels of gene expression variation [54, 55], but this has not been demonstrated for multicellular organisms, and it is not clear why levels of variability between cells should correlate with levels of variability measured at a tissue level between individuals. Given this, it is intriguing that studies at the single-cell level [5, 52] and at the tissue level [18, 53] have found similar patterns in the functions of genes with different levels of expression variability, which are also consistent with the GO categories enriched among differentially dispersed genes in this study. This is an active area of research, and there is clearly more work to be done to elucidate the sources and significance of differences in variability at the cell-to-cell and individual-to-individual levels. In both cases, care must be taken to distinguish between differences in variability arising as an artifact of hidden heterogeneity and differences in variability within a homogeneous population.

While there is a clear interest in elucidating the roles that different patterns of disruption of normal expression play, in terms of identifying cancer-related genes, any difference in the distribution of expression values between groups is of interest. Differential distribution has also been shown to provide improved feature selection for cancer classification compared to differential expression or variability alone [17]. Differential distribution analysis may therefore be preferable to separate tests of differential expression and variability both in terms of identifying potential cancer-related genes, and for studies into diagnostic or prognostic prediction.

### Identifying differentially variable or distributed genes from RNA-seq data

The hierarchical model presented here is similar to that used by BASiCS, but applied to bulk RNA-seq rather than single-cell, without the need for spike-in genes or technical replicates, and using a fully hierarchical model, where the parameters for the priors on the mean and variance parameters are not fixed, but are also estimated as part of the model. The model is further extended into a mixture model, allowing for an overall test of differential distribution in addition to separate tests for differential expression and differential dispersion.

In tests on simulated data and on real data modified to artificially induce changes in expression, tests for differential dispersion using the hierarchical model outperformed the differential dispersion test of MDSeq. While MDSeq uses a different formulation of the NB distribution, a difference in dispersion under the form used here still equates to a difference in dispersion under the MDSeq model. Unlike the hierarchical model, and most differential expression methods, MDSeq does not share information between genes, instead treating each gene independently. This may partly explain the improvement of the hierarchical model over MDSeq for differential dispersion detection. However, GAMLSS also outperformed MDSeq, and gave nearly identical performance to the hierarchical model on larger sample sizes. Which method is preferred for differential dispersion analysis may depend on the study design. In particular, the hierarchical model allows for testing at a minimum LFC, which GAMLSS does not.

In addition to the hierarchical mixture model, we considered a “hybrid” test for differential distribution that combines separate tests for differential expression and differential dispersion. While both methods were successful in identifying differentially distributed genes, false discovery curves (Figure 1C) show that, for a given number of discoveries, combining separate tests generally resulted in fewer false discoveries. This may at least in part reflect a general issue with inference from MCMC models, where the precision of parameter estimates is limited by the posterior sample size.

This limitation of posterior sampling-based inference is evident in the false discovery curves for both the differential dispersion and differential distribution tests from the hierarchical model, for which there are sometimes several hundred top-ranked genes which cannot be separated. This issue could be avoided if a suitable hierarchical model could be identified for which analytical solutions could be found for the posterior parameter distributions. This is likely to mean moving away from the NB model for RNA-seq count distributions, but the success of limma–voom in identifying differential expression using a method that focuses on appropriately modelling the mean-variance relationship rather than specifying the exact distribution of the counts [36] suggests that such an approach is worth exploring. Another way of avoiding the issue of returning many equally-ranked genes may be to incorporate information on the magnitude of changes in mean or dispersion into gene rankings. The issue may also be mitigated somewhat by performing tests at a minimum LFC, as is evident from Additional file 1: Figure S5.

Relatively small sample sizes were used in this study – up to 50 samples per group. The results show that the hierarchical model has advantages over methods that treat each gene independently for small sample sizes, but this is not enough to completely mitigate the difficulty inherent in testing for changes in variability among small datasets. While differential expression analysis can be informative even with very small samples, analysis of differential variability or distribution may be best reserved for situations where larger numbers of samples are available.

## Methods

### Hierarchical model to detect differential distribution in RNA-seq data

We assume that the observed read count *Y_ij_* for gene *j* in sample *i* follows a negative binomial distribution, parametrised by a gene-specific mean *μ_j_* and dispersion *ϕ_j_*:

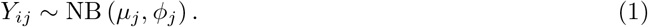

We specify log-normal priors for the mean and dispersion parameters:

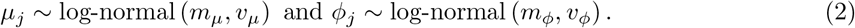

The top level of the hierarchical model consists of hyperpriors on the prior parameters for the means and dispersions. We specify normal hyperpriors for the location parameters *m_μ_* and *m_ϕ_*, and gamma hyperpriors for the scale parameters *υ_μ_* and *υ_ϕ_*. The hyperpriors were chosen to place most density on the regions of the mean and dispersion parameter distributions that are most likely to be observed in real data, but with enough density outside of these regions so as to not overly restrict posterior inference. All priors are assumed to be independent, and an adaptive MCMC sampling scheme is used to obtain posterior samples of the mean and dispersion parameters, which are the basis for inference from the hierarchical model. While posterior parameter estimates are not used directly in inference of differential expression, dispersion or distribution, parameter estimates – particularly estimates of dispersion – can be obtained as the mean of the posterior sample for each parameter. Full details of the MCMC algorithm are given in Additional file 1: Section S5.

#### Tests for differential expression and dispersion

The posterior samples resulting from the MCMC algorithm are used to form tests for differences in mean and dispersion between two groups, *A* and *B*. Given posterior samples for the mean for gene *j* in each group, *μ_jA_* and *μ_jB_*, we obtain a posterior sample of the LFC between groups by taking log_2_ *μ_jA_* — log_2_ *μ_jB_* for each MCMC iteration, and similarly for dispersion. Tail probabilities from these samples are taken as a measure of the probability that the true difference in mean or dispersion, respectively, between the two groups is not equal to zero or a given minimum LFC. Tail probabilities are obtained by constructing highest posterior density (HPD) intervals – the narrowest range of values that contains a given amount of posterior density – and iteratively finding the amount of density that uniquely defines the narrowest region that does not contain zero. Subtracting these probabilities from one gives a posterior estimate of the probability that these is no difference in mean or dispersion between groups. For tests at a minimum LFC of *c*, the narrowest region of posterior density that does not include the range [—*c, c*] is used instead of zero. Although the tail probabilities are not *p*-values, it was found that applying the Benjamini–Hochberg false discovery rate procedure [56] worked well to control the FDR in simulated data, and so this method was used where a binary decision on differential expression or dispersion was desired.

#### Mixture model for differential distribution

To detect differences in distribution between two groups, the hierarchical model is embedded in a mixture model indexed by a parameter *Z_j_*, which indicates which of the two mixture components the data for gene *j* comes from: *Z_j_* = 0 if the mean and dispersion for gene *j* are the same for both groups, and *Z_j_* = 1 if either the mean or dispersion for gene *j* differs between the groups, i.e. if there is differential distribution for that gene.

The mixture model is defined by the following model for the distribution of read counts:

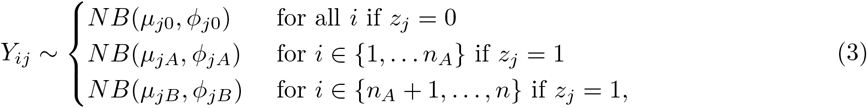

where *n* is the total number of samples and *n_A_* the number of samples in group *A*.

We assign a Bernoulli prior on the *Z_j_* with a parameter λ representing the probability that *Z_j_* = 1, that is, the probability of differential distribution. We assign a uniform prior on λ over the range (0,1). The posterior mean of λ is taken as an estimate of the proportion of differentially distributed genes, and the posterior means of the *Z_j_* are taken as estimates of the probability of differential distribution for each gene. These probabilities are used to rank genes by the strength of evidence for differential distribution. A binary decision can be made for each gene by using the posterior estimate of the proportion of differentially distributed genes to set a threshold such that the appropriate number of positive calls are made, or alternatively by using the Bayesian FDR (BFDR) [57, 58].

The R code used to implement the hierarchical model, including the MCMC algorithm and differential expression, dispersion, and distribution tests, is included in Additional file 3. The hierarchical model is implemented in an R package, DiffDist, available at https://github.com/aedanr/DiffDist.

### Datasets

#### Simulated data

Data was simulated using the compcodeR R [59]/Bioconductor [60] package [61]. Fifty simulated datasets were generated for each of 2, 5, 10, 20 and 50 samples per group, with differences in mean only for 5% of genes, dispersion only for 5%, and both mean and dispersion for 5%. Data was simulated for 20,000 genes, using the default compcodeR settings except that a minimum counts per million filter of of 0.5 was applied, and for differentially expressed genes, half were upregulated in the second group and half downregulated. compcodeR samples means and dispersions from estimates obtained from two real datasets [62, 63], and, for differentially expressed genes, multiplies or divides means by a factor of 1.5 + *x*, where *x* a random sample from an exponential distribution with mean 1. The minimum factor is therefore 1.5, and the mean factor is 2.5. To simulate differential dispersion, a dataset with no differential expression and no filtering was first generated, from which dispersion values were extracted and used as baseline dispersions. Differential dispersions were then generated using the same model as for differential expression, and data was simulated specifying these two sets of dispersions.

#### Artificially introducing differential distributions in GTEx data

Whole blood and skeletal muscle RNA-seq data from GTEx, processed by the recount2 project [64], were downloaded from https://jhubiostatistics.shinyapps.io/recount/, and gene-level counts obtained using the recount R/Bioconductor package. Samples used were limited to those from PAXGene-extracted RNA with RNA integrity number ≥ 6.9, which left 405 whole blood samples and 401 skeletal muscle samples. For each tissue type, samples were randomly selected to create ten datasets with each of 2, 5, 10, 20 and 50 samples per group. Samples were chosen without replacement for 2, 5, 10 and 20 samples per group, so that each of the ten datasets comprised different samples, and with replacement for 50 samples per group since there were insufficient samples to create ten completely distinct datasets. Counts were then adjusted in one group to introduce changes in mean and dispersion as estimated by the method-of-moments estimators. Full details are given in Additional file 1: Section S6.

#### TCGA data

RNA-seq data from TCGA were downloaded and processed as for the GTEx data. Matching pairs of primary tumour and solid tissue normal samples were retained. Where there were multiple primary tumour samples for a patient, the sample with the highest mapped read count was selected. Cancer types with at least 40 tumour–normal pairs for which lists of related genes were available from KEGG were selected, and where there were multiple histological diagnoses for a cancer type, only the most common was used: infiltrating ductal breast carcinoma; classical/usual papillary thyroid carcinoma; lung adenocarcinoma not otherwise specified (NOS); hepatocellular carcinoma; lung squamous cell carcinoma NOS; and colon adenocarcinoma. The resulting sample sizes and references for original publications are summarised in Table 1.

#### Cancer-related genes

Lists of genes previously identified as being related to each of the eight cancer types were compiled by combining lists of related genes from CGC [39], DisGeNET [40], IntOGen [41], KEGG and MalaCards [43]. Full details are given in Additional file 1: Section S7. The lists of genes obtained from each database are given in Additional file 4, and the compiled lists of cancer-related genes in Additional file 5.

### Comparative assessment of model performance

Differential dispersion performance using HM was compared against MDSeq and GAMLSS, and differential expression performance was compared against edgeR [33, 65, 66], DESeq2 [34], limma-voom [36, 67], baySeq [30] and MDSeq. The default settings were used for each method except that any gene filtering that was applied by default was not used as filtering was applied during simulation. Three versions of edgeR were initially tested: quasi-likelihood, likelihood ratio test and exact test. Quasi-likelihood gave the best results, and so only results from this version are reported. Preliminary testing showed that results were nearly identical using edgeR’s TMM normalisation [68] and DESeq2’s normalisation. TMM was used for all subsequent analyses.

The performance of the HMM differential distribution test was compared against diffVar [37, 38] and a naive method combining the results from separate differential expression and differential dispersion tests. Differential expression was assessed using edgeR for the simulated data and limma–voom for the GTEx data as these were the best-performing tests in each situation, and differential dispersion using HM. Results from the two tests were combined by taking the smaller of the *p*-value or posterior probability of no difference between groups for each gene. The resulting values were adjusted for multiple comparisons using the Benjamini–Hochberg procedure.

The R code used for each method is included in Additional file 6, and can also be found at https://github.com/aedanr/DiffDist.

### Analysis of tumour–normal comparisons

Wilcoxon rank-sum tests were used to test the ability of differential expression and differential dispersion analyses to identify cancer-related genes. For each method, genes were ranked by the strength of evidence for differential expression or dispersion, and genes previously identified as being related to each cancer type were considered as positive, and all other genes as negative.

Spearman (rank) correlation was used to assess the similarity between lists of genes identified by differential expression and differential dispersion. For a list of genes for each method defined by a *p*-value or posterior probability threshold, the Spearman correlation between the union of genes in the two lists was calculated, and a corresponding one-sided correlation hypothesis test performed, with the alternative hypothesis that the correlation was negative.

Gene set enrichment analysis was performed using the GSEA software [69], version 4.0.3, using the Human_ENSEMBL_Gene_MSigDB.v7.1 chip annotation and with genes ranked by the absolute LFC in mean or dispersion multiplied by – log_10_ *p*, where *p* is the *p*-value or posterior probability of no difference between groups. This means that both the strength of evidence for a difference between normal and tumour and the magnitude of the difference contribute to a gene’s ranking, and absolute values are used since a change in either direction is relevant. Analyses were carried out for GO terms in each of the three ontologies, with terms with less than two or more than six levels of parent terms excluded in order to avoid terms that were too general or too specific to be meaningfully interpreted. GO terms were taken from the org.Hs.eg.db annotation, and relationships between terms were obtained using the GO.db R/Bioconductor package. The resulting terms were then matched to the list of terms for each ontology in Molecular Signatures Database gene sets using the GSEABase package. To further aid the interpretability of the results, semantic similarity matrices for terms in the results lists were identified using the GOSemSim R/Bioconductor package [70], and terms with high redundancy removed using the rrvgo package, which is based on REVIGO [71], with a threshold of 0.5.

## Supporting information

Additional file 2: GSEA results

Additional file 3: MCMC algorithm

Additional file 4: Methods comparisons

Additional file 5: Cancer-related gene sources

Additional file 6: Cancer-related genes

## Acknowledgements

The authors would like to thank Thomas Lysaght and George Mundackal for their contributions to optimising the MCMC code. The Genotype–Tissue Expression (GTEx) Project was supported by the Common Fund of the Office of the Director of the National Institutes of Health, USA, and by NCI, NHGRI, NHLBI, NIDA, NIMH and NINDS. The results on tumour—normal comparisons are based upon data generated by the TCGA Research Network: https://www.cancer.gov.tcga. The authors would like to thank the anonymous specimen donors for their essential contribution to this research.

## Abbreviations

BFDR: Bayesian false discovery rate
CGC: Cancer Genome Census
CV: Coefficient of variation
FDR: False discovery rate
GAMLSS: Generalised additive models for location, shape and scale
GO: Gene ontology
GTEx: Genotype-Tissue Expression
HM: Hierarchical model
HMM: Hierarchical mixture model
HPD: Highest posterior density
KEGG: Kyoto Encyclopedia of Genes and Genomes
LFC: Log fold change
MCMC: Markov chain Monte Carlo
NB: Negative binomial
NOS: Not otherwise specified
RNA-seq: RNA sequencing
TCGA: The Cancer Genome Atlas

## Availability of data and materials

The code for the hierarchical model and associated tests is included as Additional file 3, and the model is implemented in an R package, DiffDist, available at https://github.com/aedanr/DiffDist [72]. The code used to run each of the differential expression, dispersion and distribution methods for the methods comparisons is included as Additional file 4.

TCGA and GTEx data as processed by the recount2 project [73] were downloaded from https://jhubiostatistics.shinyapps.io/recount/. GSEA software was downloaded from https://www.gsea-msigdb.org/gsea/index.jsp. Additional file 2 gives the GSEA results for the top 10 terms identified in each ontology for each cancer type for differential expression analysis using limma-voom and differential variability using the hierarchical model and GAMLSS. Full details of the process for obtaining lists of genes previously identified as being related to each cancer are given in Additional file 1, Section S7. The gene lists obtained from each database are given in Additional file 4, and the compiled lists of cancer-related genes in Additional file 5.

## Ethics approval and consent to participate

Not applicable.

## Consent for publication

Not applicable.

## Authors’ contributions

AGKR, PJK and DRC conceived the study. AGKR developed and implemented the hierarchical model with input from PJK. AGKR designed and performed the analysis, interpreted the results and wrote the manuscript with input from PJK and DRC. All authors read and approved the final manuscript.

## Additional files

### Additional file 1 — Supplementary results and methods (.pdf)

Section S1 provides the results of comparative analysis of differential expression tests using the hierarchical model and existing methods. Section S2 shows the results of differential dispersion comparisons with a wider range of sample sizes. Section S3 shows the results of differential distribution comparisons with a wider range of sample sizes. Section S4 provides the results of the investigation into differential dispersion and distribution on all eight cancer types analysed. Section S5 contains additional details on the hierarchical model, the MCMC algorithm used to generate posterior samples, and the test for differential distribution using the hierarchical mixture model. Section S6 gives further details on introducing changes in mean and dispersion in the GTEx datasets. Section S7 details the process of collating lists of previously-identified cancer-related genes.

### Additional file 2 — GSEA results(.xlsx)

Lists of the 10 most significantly enriched GO terms in each ontology for each of the eight cancer types from differential expression analysis using limma–voom and differential variability using the hierarchical model and GAMLSS.

### Additional file 3 — MCMC algorithm (.pdf)

R code for the MCMC algorithm.

### Additional file 4 — Methods comparisons (.pdf)

R code for methods comparisons.

### Additional file 5 — Cancer-related gene sources (.xlsx)

Lists of genes obtained from CGC, DisGeNET, IntOGen, KEGG and MalaCards.

### Additional file 6 — Cancer-related genes (.xlsx)

Compiled lists of cancer-related genes.

