## Additional file 3: MCMC algorithm for "Identification of differentially distributed gene expression and distinct sets of cancer-related genes identified by changes in mean and variability"

### Additional file 2 for *Identification of differentially distributed gene expression and distinct sets of cancer-related genes identified by changes in mean and variability*: R code for the MCMC algorithm and tests for differential expression, dispersion and distribution

#### Top-level function to run adaptive MCMC algorithm

```
ln_hmm_adapt_3_chains <- function(counts, groups, chain.length=2000,
                                  initial.chain.length=100,
                                  adapt.chain.length=100) {

  require(coda)
  genes <- ncol(counts)
  counts1 <- counts[groups==1,]
  counts2 <- counts[groups==2,]
  samples0 <- nrow(counts)
  samples1 <- nrow(counts1)
  samples2 <- nrow(counts2)
  sample.means0 <- pmax(colMeans(counts), 0.01)
  sample.means1 <- pmax(colMeans(counts1), 0.01)
  sample.means2 <- pmax(colMeans(counts2), 0.01)
  sample.vars0 <- apply(counts, 2, var)
  sample.vars1 <- apply(counts1, 2, var)
  sample.vars2 <- apply(counts2, 2, var)
  rm(counts1, counts2)
  sample.disps0 <- pmax(sample.vars0-sample.means0, 0.01) / pmax(sample.means0^2, 0.1)
  sample.disps1 <- pmax(sample.vars1-sample.means1, 0.01) / pmax(sample.means1^2, 0.1)
  sample.disps2 <- pmax(sample.vars2-sample.means2, 0.01) / pmax(sample.means2^2, 0.1)
  rm(sample.vars0, sample.vars1, sample.vars2)

  # Set proposal scales
  message("initialising...")
  adapt.mcmc <- ln_hmm_1_chain(
    counts=counts, groups=groups, chain.length=initial.chain.length, thin=1,
    inits=list("means0"=sample.means0, "means1"=sample.means1, "means2"=sample.means2,
               "disps0"=sample.disps0, "disps1"=sample.disps1, "disps2"=sample.disps2,
               "mean.prior.location"=1, "disp.prior.location"=1,
               "mean.prior.scale"=1, "disp.prior.scale"=1),
    mean.proposal.scales0=rep(0.2, ncol(counts)),
    mean.proposal.scales1=rep(0.2, ncol(counts)),
    mean.proposal.scales2=rep(0.2, ncol(counts)),
    disp.proposal.scales0=rep(0.5, ncol(counts)),
    disp.proposal.scales1=rep(0.5, ncol(counts)),
    disp.proposal.scales2=rep(0.5, ncol(counts)),
    mean.prior.scale.proposal.sd=0.1, disp.prior.scale.proposal.sd=0.4
  )
  rm(sample.means0, sample.means1, sample.means2,
     sample.disps0, sample.disps1, sample.disps2)

  inits1.adapt <- list(
    "means0"=apply(adapt.mcmc$posterior.means0, 2, min)/5,
```

```
  "means1"=apply(adapt.mcmc$posterior.means1, 2, min)/5,
  "means2"=apply(adapt.mcmc$posterior.means2, 2, min)/5,
  "disps0"=apply(adapt.mcmc$posterior.disps0, 2, min)/10,
  "disps1"=apply(adapt.mcmc$posterior.disps1, 2, min)/10,
  "disps2"=apply(adapt.mcmc$posterior.disps2, 2, min)/10,
  "mean.prior.location"=min(adapt.mcmc$posterior.mean.prior.location)/2,
  "mean.prior.scale"=min(adapt.mcmc$posterior.mean.prior.scale)/2,
  "disp.prior.location"=min(adapt.mcmc$posterior.disp.prior.location)/2,
  "disp.prior.scale"=min(adapt.mcmc$posterior.disp.prior.scale)/2
)
inits2.adapt <- list(
  "means0"=colMeans(adapt.mcmc$posterior.means0),
  "means1"=colMeans(adapt.mcmc$posterior.means1),
  "means2"=colMeans(adapt.mcmc$posterior.means2),
  "disps0"=colMeans(adapt.mcmc$posterior.disps0),
  "disps1"=colMeans(adapt.mcmc$posterior.disps1),
  "disps2"=colMeans(adapt.mcmc$posterior.disps2),
  "mean.prior.location"=mean(adapt.mcmc$posterior.mean.prior.location),
  "mean.prior.scale"=mean(adapt.mcmc$posterior.mean.prior.scale),
  "disp.prior.location"=mean(adapt.mcmc$posterior.disp.prior.location),
  "disp.prior.scale"=mean(adapt.mcmc$posterior.disp.prior.scale)
)
inits3.adapt <- list(
  "means0"=apply(adapt.mcmc$posterior.means0, 2, max)*5,
  "means1"=apply(adapt.mcmc$posterior.means1, 2, max)*5,
  "means2"=apply(adapt.mcmc$posterior.means2, 2, max)*5,
  "disps0"=apply(adapt.mcmc$posterior.disps0, 2, max)*10,
  "disps1"=apply(adapt.mcmc$posterior.disps1, 2, max)*10,
  "disps2"=apply(adapt.mcmc$posterior.disps2, 2, max)*10,
  "mean.prior.location"=max(adapt.mcmc$posterior.mean.prior.location)*2,
  "mean.prior.scale"=max(adapt.mcmc$posterior.mean.prior.scale)*2,
  "disp.prior.location"=max(adapt.mcmc$posterior.disp.prior.location)*2,
  "disp.prior.scale"=max(adapt.mcmc$posterior.disp.prior.scale)*2
)

message("optimising proposal distributions...")
current.mean.proposal.scales0 <- adapt.mcmc$mean.proposal.scales0 *
  (1 + 2.25 * (adapt.mcmc$accept.means0 - 0.44))
current.mean.proposal.scales1 <- adapt.mcmc$mean.proposal.scales1 *
  (1 + 2.25 * (adapt.mcmc$accept.means1 - 0.44))
current.mean.proposal.scales2 <- adapt.mcmc$mean.proposal.scales2 *
  (1 + 2.25 * (adapt.mcmc$accept.means2 - 0.44))
current.disp.proposal.scales0 <- adapt.mcmc$disp.proposal.scales0 *
  (1 + 2.25 * (adapt.mcmc$accept.disps0 - 0.44))
current.disp.proposal.scales1 <- adapt.mcmc$disp.proposal.scales1 *
  (1 + 2.25 * (adapt.mcmc$accept.disps1 - 0.44))
current.disp.proposal.scales2 <- adapt.mcmc$disp.proposal.scales2 *
  (1 + 2.25 * (adapt.mcmc$accept.disps2 - 0.44))
current.mean.prior.scale.proposal.sd <- adapt.mcmc$mean.prior.scale.proposal.sd *
  (1 + 2.25 * (adapt.mcmc$accept.mean.prior.scale - 0.44))
current.disp.prior.scale.proposal.sd <- adapt.mcmc$disp.prior.scale.proposal.sd *
  (1 + 2.25 * (adapt.mcmc$accept.disp.prior.scale - 0.44))
```

```
adapt.mcmc <- ln_hmm_3_chains(
  counts=counts, groups=groups, chain.length=adapt.chain.length,
  thin=adapt.chain.length,
  inits1=inits1.adapt, inits2=inits2.adapt, inits3=inits3.adapt,
  mean.proposal.scales0=current.mean.proposal.scales0,
  mean.proposal.scales1=current.mean.proposal.scales1,
  mean.proposal.scales2=current.mean.proposal.scales2,
  disp.proposal.scales0=current.disp.proposal.scales0,
  disp.proposal.scales1=current.disp.proposal.scales1,
  disp.proposal.scales2=current.disp.proposal.scales2,
  mean.prior.scale.proposal.sd=current.mean.prior.scale.proposal.sd,
  disp.prior.scale.proposal.sd=current.disp.prior.scale.proposal.sd
)
rm(inits1.adapt, inits2.adapt, inits3.adapt)

current.means0.1 <- adapt.mcmc$posterior.means0.1
current.means1.1 <- adapt.mcmc$posterior.means1.1
current.means2.1 <- adapt.mcmc$posterior.means2.1
current.means0.2 <- adapt.mcmc$posterior.means0.2
current.means1.2 <- adapt.mcmc$posterior.means1.2
current.means2.2 <- adapt.mcmc$posterior.means2.2
current.means0.3 <- adapt.mcmc$posterior.means0.3
current.means1.3 <- adapt.mcmc$posterior.means1.3
current.means2.3 <- adapt.mcmc$posterior.means2.3
current.disps0.1 <- adapt.mcmc$posterior.disps0.1
current.disps1.1 <- adapt.mcmc$posterior.disps1.1
current.disps2.1 <- adapt.mcmc$posterior.disps2.1
current.disps0.2 <- adapt.mcmc$posterior.disps0.2
current.disps1.2 <- adapt.mcmc$posterior.disps1.2
current.disps2.2 <- adapt.mcmc$posterior.disps2.2
current.disps0.3 <- adapt.mcmc$posterior.disps0.3
current.disps1.3 <- adapt.mcmc$posterior.disps1.3
current.disps2.3 <- adapt.mcmc$posterior.disps2.3
current.mean.prior.location.1 <- adapt.mcmc$posterior.mean.prior.location.1
current.mean.prior.location.2 <- adapt.mcmc$posterior.mean.prior.location.2
current.mean.prior.location.3 <- adapt.mcmc$posterior.mean.prior.location.3
current.mean.prior.scale.1 <- adapt.mcmc$posterior.mean.prior.scale.1
current.mean.prior.scale.2 <- adapt.mcmc$posterior.mean.prior.scale.2
current.mean.prior.scale.3 <- adapt.mcmc$posterior.mean.prior.scale.3
current.disp.prior.location.1 <- adapt.mcmc$posterior.disp.prior.location.1
current.disp.prior.location.2 <- adapt.mcmc$posterior.disp.prior.location.2
current.disp.prior.location.3 <- adapt.mcmc$posterior.disp.prior.location.3
current.disp.prior.scale.1 <- adapt.mcmc$posterior.disp.prior.scale.1
current.disp.prior.scale.2 <- adapt.mcmc$posterior.disp.prior.scale.2
current.disp.prior.scale.3 <- adapt.mcmc$posterior.disp.prior.scale.3

accept.means0 <- (adapt.mcmc$accept.means0.1 + adapt.mcmc$accept.means0.2 +
                    adapt.mcmc$accept.means0.3)/3
accept.means1 <- (adapt.mcmc$accept.means1.1 + adapt.mcmc$accept.means1.2 +
                    adapt.mcmc$accept.means1.3)/3
accept.means2 <- (adapt.mcmc$accept.means2.1 + adapt.mcmc$accept.means2.2 +
                    adapt.mcmc$accept.means2.3)/3
accept.disps0 <- (adapt.mcmc$accept.disps0.1 + adapt.mcmc$accept.disps0.2 +
```

```
                     adapt.mcmc$accept.disps0.3)/3
accept.disps1 <- (adapt.mcmc$accept.disps1.1 + adapt.mcmc$accept.disps1.2 +
                     adapt.mcmc$accept.disps1.3)/3
accept.disps2 <- (adapt.mcmc$accept.disps2.1 + adapt.mcmc$accept.disps2.2 +
                     adapt.mcmc$accept.disps2.3)/3
accept.mean.prior.scale <- mean(c(adapt.mcmc$accept.mean.prior.scale.1,
                                   adapt.mcmc$accept.mean.prior.scale.2,
                                   adapt.mcmc$accept.mean.prior.scale.3))
accept.disp.prior.scale <- mean(c(adapt.mcmc$accept.disp.prior.scale.1,
                                   adapt.mcmc$accept.disp.prior.scale.2,
                                   adapt.mcmc$accept.disp.prior.scale.3))
adapt.runs <- 1

while (sum(accept.means0<0.24 | accept.means0>0.64 |
           accept.means1<0.24 | accept.means1>0.64 |
           accept.means2<0.24 | accept.means2>0.64 |
           accept.disps0<0.24 | accept.disps0>0.64 |
           accept.disps1<0.24 | accept.disps1>0.64 |
           accept.disps2<0.24 | accept.disps2>0.64 |
           accept.mean.prior.scale<0.24 | accept.mean.prior.scale>0.64 |
           accept.disp.prior.scale<0.24 | accept.disp.prior.scale>0.64) > 0) {
  current.mean.proposal.scales0 <-
    current.mean.proposal.scales0 * (1 + 2.25 * (accept.means0-0.44))
  current.mean.proposal.scales1 <-
    current.mean.proposal.scales1 * (1 + 2.25 * (accept.means1-0.44))
  current.mean.proposal.scales2 <-
    current.mean.proposal.scales2 * (1 + 2.25 * (accept.means2-0.44))
  current.disp.proposal.scales0 <-
    current.disp.proposal.scales0 * (1 + 2.25 * (accept.disps0-0.44))
  current.disp.proposal.scales1 <-
    current.disp.proposal.scales1 * (1 + 2.25 * (accept.disps1-0.44))
  current.disp.proposal.scales2 <-
    current.disp.proposal.scales2 * (1 + 2.25 * (accept.disps2-0.44))
  current.mean.prior.scale.proposal.sd <- current.mean.prior.scale.proposal.sd *
    (1 + 2.25 * (accept.mean.prior.scale-0.44))
  current.disp.prior.scale.proposal.sd <- current.disp.prior.scale.proposal.sd *
    (1 + 2.25 * (accept.disp.prior.scale-0.44))

  adapt.mcmc <- ln_hmm_3_chains(
    counts=counts, groups=groups, chain.length=adapt.chain.length,
    thin=adapt.chain.length,
    inits1=list("means0"=current.means0.1, "means1"=current.means1.1,
                "means2"=current.means2.1, "disps0"=current.disps0.1,
                "disps1"=current.disps1.1, "disps2"=current.disps2.1,
                "mean.prior.location"=current.mean.prior.location.1,
                "mean.prior.scale"=current.mean.prior.scale.1,
                "disp.prior.location"=current.disp.prior.location.1,
                "disp.prior.scale"=current.disp.prior.scale.1),
    inits2=list("means0"=current.means0.2, "means1"=current.means1.2,
                "means2"=current.means2.2, "disps0"=current.disps0.2,
                "disps1"=current.disps1.2, "disps2"=current.disps2.2,
                "mean.prior.location"=current.mean.prior.location.2,
                "mean.prior.scale"=current.mean.prior.scale.2,
```

```
                "disp.prior.location"=current.disp.prior.location.2,
                "disp.prior.scale"=current.disp.prior.scale.2),
  inits3=list("means0"=current.means0.3, "means1"=current.means1.3,
                "means2"=current.means2.3, "disps0"=current.disps0.3,
                "disps1"=current.disps1.3, "disps2"=current.disps2.3,
                "mean.prior.location"=current.mean.prior.location.3,
                "mean.prior.scale"=current.mean.prior.scale.3,
                "disp.prior.location"=current.disp.prior.location.3,
                "disp.prior.scale"=current.disp.prior.scale.3),
  mean.proposal.scales0=current.mean.proposal.scales0,
  mean.proposal.scales1=current.mean.proposal.scales1,
  mean.proposal.scales2=current.mean.proposal.scales2,
  disp.proposal.scales0=current.disp.proposal.scales0,
  disp.proposal.scales1=current.disp.proposal.scales1,
  disp.proposal.scales2=current.disp.proposal.scales2,
  mean.prior.scale.proposal.sd=current.mean.prior.scale.proposal.sd,
  disp.prior.scale.proposal.sd=current.disp.prior.scale.proposal.sd
)

current.means0.1 <- adapt.mcmc$posterior.means0.1
current.means1.1 <- adapt.mcmc$posterior.means1.1
current.means2.1 <- adapt.mcmc$posterior.means2.1
current.means0.2 <- adapt.mcmc$posterior.means0.2
current.means1.2 <- adapt.mcmc$posterior.means1.2
current.means2.2 <- adapt.mcmc$posterior.means2.2
current.means0.3 <- adapt.mcmc$posterior.means0.3
current.means1.3 <- adapt.mcmc$posterior.means1.3
current.means2.3 <- adapt.mcmc$posterior.means2.3
current.disps0.1 <- adapt.mcmc$posterior.disps0.1
current.disps1.1 <- adapt.mcmc$posterior.disps1.1
current.disps2.1 <- adapt.mcmc$posterior.disps2.1
current.disps0.2 <- adapt.mcmc$posterior.disps0.2
current.disps1.2 <- adapt.mcmc$posterior.disps1.2
current.disps2.2 <- adapt.mcmc$posterior.disps2.2
current.disps0.3 <- adapt.mcmc$posterior.disps0.3
current.disps1.3 <- adapt.mcmc$posterior.disps1.3
current.disps2.3 <- adapt.mcmc$posterior.disps2.3
current.mean.prior.location.1 <- adapt.mcmc$posterior.mean.prior.location.1
current.mean.prior.location.2 <- adapt.mcmc$posterior.mean.prior.location.2
current.mean.prior.location.3 <- adapt.mcmc$posterior.mean.prior.location.3
current.mean.prior.scale.1 <- adapt.mcmc$posterior.mean.prior.scale.1
current.mean.prior.scale.2 <- adapt.mcmc$posterior.mean.prior.scale.2
current.mean.prior.scale.3 <- adapt.mcmc$posterior.mean.prior.scale.3
current.disp.prior.location.1 <- adapt.mcmc$posterior.disp.prior.location.1
current.disp.prior.location.2 <- adapt.mcmc$posterior.disp.prior.location.2
current.disp.prior.location.3 <- adapt.mcmc$posterior.disp.prior.location.3
current.disp.prior.scale.1 <- adapt.mcmc$posterior.disp.prior.scale.1
current.disp.prior.scale.2 <- adapt.mcmc$posterior.disp.prior.scale.2
current.disp.prior.scale.3 <- adapt.mcmc$posterior.disp.prior.scale.3

accept.means0 <- (adapt.mcmc$accept.means0.1 + adapt.mcmc$accept.means0.2 +
                    adapt.mcmc$accept.means0.3)/3
accept.means1 <- (adapt.mcmc$accept.means1.1 + adapt.mcmc$accept.means1.2 +
```

```
                      adapt.mcmc$accept.means1.3)/3
  accept.means2 <- (adapt.mcmc$accept.means2.1 + adapt.mcmc$accept.means2.2 +
                      adapt.mcmc$accept.means2.3)/3
  accept.disps0 <- (adapt.mcmc$accept.disps0.1 + adapt.mcmc$accept.disps0.2 +
                      adapt.mcmc$accept.disps0.3)/3
  accept.disps1 <- (adapt.mcmc$accept.disps1.1 + adapt.mcmc$accept.disps1.2 +
                      adapt.mcmc$accept.disps1.3)/3
  accept.disps2 <- (adapt.mcmc$accept.disps2.1 + adapt.mcmc$accept.disps2.2 +
                      adapt.mcmc$accept.disps2.3)/3
  accept.mean.prior.scale <- mean(c(adapt.mcmc$accept.mean.prior.scale.1,
                                    adapt.mcmc$accept.mean.prior.scale.2,
                                    adapt.mcmc$accept.mean.prior.scale.3))
  accept.disp.prior.scale <- mean(c(adapt.mcmc$accept.disp.prior.scale.1,
                                    adapt.mcmc$accept.disp.prior.scale.2,
                                    adapt.mcmc$accept.disp.prior.scale.3))

  adapt.runs <- adapt.runs+1
}
rm(adapt.mcmc)

inits1 <- list("means0"=current.means0.1, "means1"=current.means1.1,
               "means2"=current.means2.1, "disps0"=current.disps0.1,
               "disps1"=current.disps1.1, "disps2"=current.disps2.1,
               "mean.prior.location"=current.mean.prior.location.1,
               "mean.prior.scale"=current.mean.prior.scale.1,
               "disp.prior.location"=current.disp.prior.location.1,
               "disp.prior.scale"=current.disp.prior.scale.1)
inits2 <- list("means0"=current.means0.2, "means1"=current.means1.2,
               "means2"=current.means2.2, "disps0"=current.disps0.2,
               "disps1"=current.disps1.2, "disps2"=current.disps2.2,
               "mean.prior.location"=current.mean.prior.location.2,
               "mean.prior.scale"=current.mean.prior.scale.2,
               "disp.prior.location"=current.disp.prior.location.2,
               "disp.prior.scale"=current.disp.prior.scale.2)
inits3 <- list("means0"=current.means0.3, "means1"=current.means1.3,
               "means2"=current.means2.3, "disps0"=current.disps0.3,
               "disps1"=current.disps1.3, "disps2"=current.disps2.3,
               "mean.prior.location"=current.mean.prior.location.3,
               "mean.prior.scale"=current.mean.prior.scale.3,
               "disp.prior.location"=current.disp.prior.location.3,
               "disp.prior.scale"=current.disp.prior.scale.3)
mean.proposal.scales0 <- current.mean.proposal.scales0
mean.proposal.scales1 <- current.mean.proposal.scales1
mean.proposal.scales2 <- current.mean.proposal.scales2
disp.proposal.scales0 <- current.disp.proposal.scales0
disp.proposal.scales1 <- current.disp.proposal.scales1
disp.proposal.scales2 <- current.disp.proposal.scales2
mean.prior.scale.proposal.sd <- current.mean.prior.scale.proposal.sd
disp.prior.scale.proposal.sd <- current.disp.prior.scale.proposal.sd

# Run MCMC ####
message("running mcmc...")
final.chains <- ln_hmm_3_chains(
```

```
  counts=counts, groups=groups, chain.length=chain.length, thin=1,
  inits1=inits1, inits2=inits2, inits3=inits3,
  mean.proposal.scales0=mean.proposal.scales0,
  mean.proposal.scales1=mean.proposal.scales1,
  mean.proposal.scales2=mean.proposal.scales2,
  disp.proposal.scales0=disp.proposal.scales0,
  disp.proposal.scales1=disp.proposal.scales1,
  disp.proposal.scales2=disp.proposal.scales2,
  mean.prior.scale.proposal.sd=mean.prior.scale.proposal.sd,
  disp.prior.scale.proposal.sd=disp.prior.scale.proposal.sd
)
rm(inits1, inits2, inits3)

means0 <- mcmc.list(mcmc(final.chains$posterior.means0.1),
                    mcmc(final.chains$posterior.means0.2),
                    mcmc(final.chains$posterior.means0.3))
means1 <- mcmc.list(mcmc(final.chains$posterior.means1.1),
                    mcmc(final.chains$posterior.means1.2),
                    mcmc(final.chains$posterior.means1.3))
means2 <- mcmc.list(mcmc(final.chains$posterior.means2.1),
                    mcmc(final.chains$posterior.means2.2),
                    mcmc(final.chains$posterior.means2.3))
disps0 <- mcmc.list(mcmc(final.chains$posterior.disps0.1),
                    mcmc(final.chains$posterior.disps0.2),
                    mcmc(final.chains$posterior.disps0.3))
disps1 <- mcmc.list(mcmc(final.chains$posterior.disps1.1),
                    mcmc(final.chains$posterior.disps1.2),
                    mcmc(final.chains$posterior.disps1.3))
disps2 <- mcmc.list(mcmc(final.chains$posterior.disps2.1),
                    mcmc(final.chains$posterior.disps2.2),
                    mcmc(final.chains$posterior.disps2.3))
mean.prior.location <-
  mcmc.list(mcmc(final.chains$posterior.mean.prior.location.1),
            mcmc(final.chains$posterior.mean.prior.location.2),
            mcmc(final.chains$posterior.mean.prior.location.3))
mean.prior.scale <-
  mcmc.list(mcmc(final.chains$posterior.mean.prior.scale.1),
            mcmc(final.chains$posterior.mean.prior.scale.2),
            mcmc(final.chains$posterior.mean.prior.scale.3))
disp.prior.location <-
  mcmc.list(mcmc(final.chains$posterior.disp.prior.location.1),
            mcmc(final.chains$posterior.disp.prior.location.2),
            mcmc(final.chains$posterior.disp.prior.location.3))
disp.prior.scale <-
  mcmc.list(mcmc(final.chains$posterior.disp.prior.scale.1),
            mcmc(final.chains$posterior.disp.prior.scale.2),
            mcmc(final.chains$posterior.disp.prior.scale.3))
indicators <- mcmc.list(mcmc(final.chains$posterior.indicators.1),
                        mcmc(final.chains$posterior.indicators.2),
                        mcmc(final.chains$posterior.indicators.3))
proportion <- mcmc.list(mcmc(final.chains$posterior.proportion.1),
                        mcmc(final.chains$posterior.proportion.2),
                        mcmc(final.chains$posterior.proportion.3))
```

```
  rm(final.chains)

  return(list(
    "chain.length"=chain.length, "adaptive.runs"=adapt.runs,
    "mean.proposal.scales0"=mean.proposal.scales0,
    "mean.proposal.scales1"=mean.proposal.scales1,
    "mean.proposal.scales2"=mean.proposal.scales2,
    "disp.proposal.scales0"=disp.proposal.scales0,
    "disp.proposal.scales1"=disp.proposal.scales1,
    "disp.proposal.scales2"=disp.proposal.scales2,
    "mean.prior.scale.proposal.sd"=mean.prior.scale.proposal.sd,
    "disp.prior.scale.proposal.sd"=disp.prior.scale.proposal.sd,
    "means0"=means0,"means1"=means1, "means2"=means2,
    "disps0"=disps0, "disps1"=disps1, "disps2"=disps2,
    "mean.prior.location"=mean.prior.location, "disp.prior.location"=disp.prior.location,
    "mean.prior.scale"=mean.prior.scale, "disp.prior.scale"=disp.prior.scale,
    "indicators"=indicators, "proportion"=proportion
    ))

}
```

##### Function to run three parallel MCMC chains

```
ln_hmm_3_chains <- function(
  counts, groups, chain.length, thin=1, inits1, inits2, inits3,
  mean.proposal.scales0=rep(0.2, ncol(counts)),
  mean.proposal.scales1=rep(0.2, ncol(counts)),
  mean.proposal.scales2=rep(0.2, ncol(counts)),
  disp.proposal.scales0=rep(0.5, ncol(counts)),
  disp.proposal.scales1=rep(0.5, ncol(counts)),
  disp.proposal.scales2=rep(0.5, ncol(counts)),
  mean.prior.scale.proposal.sd=0.1, disp.prior.scale.proposal.sd=0.4
  ) {

  genes <- ncol(counts)
  counts1 <- counts[groups==1,]
  counts2 <- counts[groups==2,]
  samples0 <- nrow(counts)
  samples1 <- nrow(counts1)
  samples2 <- nrow(counts2)
  sample.means0 <- colMeans(counts)
  sample.means1 <- colMeans(counts1)
  sample.means2 <- colMeans(counts2)
  sqrt.mean.proposal.scales0 <- sqrt(mean.proposal.scales0)
  sqrt.mean.proposal.scales1 <- sqrt(mean.proposal.scales1)
  sqrt.mean.proposal.scales2 <- sqrt(mean.proposal.scales2)
  sqrt.disp.proposal.scales0 <- sqrt(disp.proposal.scales0)
  sqrt.disp.proposal.scales1 <- sqrt(disp.proposal.scales1)
  sqrt.disp.proposal.scales2 <- sqrt(disp.proposal.scales2)

  # Create empty posterior sample matrices and vectors ####
  posterior.means0.1 <- matrix(nrow=chain.length/thin, ncol=genes)
```

```r
posterior.means1.1 <- matrix(nrow=chain.length/thin, ncol=genes)
posterior.means2.1 <- matrix(nrow=chain.length/thin, ncol=genes)
posterior.means0.2 <- matrix(nrow=chain.length/thin, ncol=genes)
posterior.means1.2 <- matrix(nrow=chain.length/thin, ncol=genes)
posterior.means2.2 <- matrix(nrow=chain.length/thin, ncol=genes)
posterior.means0.3 <- matrix(nrow=chain.length/thin, ncol=genes)
posterior.means1.3 <- matrix(nrow=chain.length/thin, ncol=genes)
posterior.means2.3 <- matrix(nrow=chain.length/thin, ncol=genes)
posterior.disps0.1 <- matrix(nrow=chain.length/thin, ncol=genes)
posterior.disps1.1 <- matrix(nrow=chain.length/thin, ncol=genes)
posterior.disps2.1 <- matrix(nrow=chain.length/thin, ncol=genes)
posterior.disps0.2 <- matrix(nrow=chain.length/thin, ncol=genes)
posterior.disps1.2 <- matrix(nrow=chain.length/thin, ncol=genes)
posterior.disps2.2 <- matrix(nrow=chain.length/thin, ncol=genes)
posterior.disps0.3 <- matrix(nrow=chain.length/thin, ncol=genes)
posterior.disps1.3 <- matrix(nrow=chain.length/thin, ncol=genes)
posterior.disps2.3 <- matrix(nrow=chain.length/thin, ncol=genes)
posterior.mean.prior.location.1 <- numeric(chain.length/thin)
posterior.mean.prior.location.2 <- numeric(chain.length/thin)
posterior.mean.prior.location.3 <- numeric(chain.length/thin)
posterior.mean.prior.scale.1 <- numeric(chain.length/thin)
posterior.mean.prior.scale.2 <- numeric(chain.length/thin)
posterior.mean.prior.scale.3 <- numeric(chain.length/thin)
posterior.disp.prior.location.1 <- numeric(chain.length/thin)
posterior.disp.prior.location.2 <- numeric(chain.length/thin)
posterior.disp.prior.location.3 <- numeric(chain.length/thin)
posterior.disp.prior.scale.1 <- numeric(chain.length/thin)
posterior.disp.prior.scale.2 <- numeric(chain.length/thin)
posterior.disp.prior.scale.3 <- numeric(chain.length/thin)
posterior.indicators.1 <- matrix(nrow=chain.length/thin, ncol=genes)
posterior.indicators.2 <- matrix(nrow=chain.length/thin, ncol=genes)
posterior.indicators.3 <- matrix(nrow=chain.length/thin, ncol=genes)
posterior.proportion.1 <- numeric(chain.length/thin)
posterior.proportion.2 <- numeric(chain.length/thin)
posterior.proportion.3 <- numeric(chain.length/thin)

# Initial values ####
current.posterior.means0.1 <- inits1$means0
current.posterior.means1.1 <- inits1$means1
current.posterior.means2.1 <- inits1$means2
current.posterior.means0.2 <- inits2$means0
current.posterior.means1.2 <- inits2$means1
current.posterior.means2.2 <- inits2$means2
current.posterior.means0.3 <- inits3$means0
current.posterior.means1.3 <- inits3$means1
current.posterior.means2.3 <- inits3$means2
current.posterior.disps0.1 <- inits1$disps0
current.posterior.disps1.1 <- inits1$disps1
current.posterior.disps2.1 <- inits1$disps2
current.posterior.disps0.2 <- inits2$disps0
current.posterior.disps1.2 <- inits2$disps1
current.posterior.disps2.2 <- inits2$disps2
current.posterior.disps0.3 <- inits3$disps0
```

```
current.posterior.disps1.3 <- inits3$disps1
current.posterior.disps2.3 <- inits3$disps2
current.posterior.mean.prior.location.1 <- inits1$mean.prior.location
current.posterior.mean.prior.location.2 <- inits2$mean.prior.location
current.posterior.mean.prior.location.3 <- inits3$mean.prior.location
current.posterior.mean.prior.scale.1 <- inits1$mean.prior.scale
current.posterior.mean.prior.scale.2 <- inits2$mean.prior.scale
current.posterior.mean.prior.scale.3 <- inits3$mean.prior.scale
current.posterior.disp.prior.location.1 <- inits1$disp.prior.location
current.posterior.disp.prior.location.2 <- inits2$disp.prior.location
current.posterior.disp.prior.location.3 <- inits3$disp.prior.location
current.posterior.disp.prior.scale.1 <- inits1$disp.prior.scale
current.posterior.disp.prior.scale.2 <- inits2$disp.prior.scale
current.posterior.disp.prior.scale.3 <- inits3$disp.prior.scale
current.posterior.indicators.1 <- numeric(genes)
current.posterior.indicators.2 <- numeric(genes)
current.posterior.indicators.3 <- numeric(genes)
current.posterior.proportion.1 = 0.5
current.posterior.proportion.2 = 0.5
current.posterior.proportion.3 = 0.5

# Create acceptance rate vectors/variables ####
accept.means0.1 <- numeric(genes)
accept.means1.1 <- numeric(genes)
accept.means2.1 <- numeric(genes)
accept.means0.2 <- numeric(genes)
accept.means1.2 <- numeric(genes)
accept.means2.2 <- numeric(genes)
accept.means0.3 <- numeric(genes)
accept.means1.3 <- numeric(genes)
accept.means2.3 <- numeric(genes)
accept.disps0.1 <- numeric(genes)
accept.disps1.1 <- numeric(genes)
accept.disps2.1 <- numeric(genes)
accept.disps0.2 <- numeric(genes)
accept.disps1.2 <- numeric(genes)
accept.disps2.2 <- numeric(genes)
accept.disps0.3 <- numeric(genes)
accept.disps1.3 <- numeric(genes)
accept.disps2.3 <- numeric(genes)
accept.mean.prior.scale.1 <- 0
accept.mean.prior.scale.2 <- 0
accept.mean.prior.scale.3 <- 0
accept.disp.prior.scale.1 <- 0
accept.disp.prior.scale.2 <- 0
accept.disp.prior.scale.3 <- 0

# Run MCMC ####
for (iter in 1:chain.length) {
  # Metropolis updates for overall per-gene means ####
  # Chain 1
  proposed.posterior.means0.1 <- rlnorm(
    n=genes, meanlog=log(current.posterior.means0.1), sdlog=sqrt.mean.proposal.scales0
```

```
)

replace <- log(runif(genes)) <=
  mean.conditional.log.posterior(
    n=samples0, sample.means=sample.means0,
    means=proposed.posterior.means0.1,
    disps=current.posterior.disps0.1,
    prior.location=current.posterior.mean.prior.location.1,
    prior.scale=current.posterior.mean.prior.scale.1
  ) +
  (log(proposed.posterior.means0.1) +
     (log(proposed.posterior.means0.1) -
        log(current.posterior.means0.1))^2 / (2*mean.proposal.scales0)) -
  mean.conditional.log.posterior(
    n=samples0, sample.means=sample.means0,
    means=current.posterior.means0.1,
    disps=current.posterior.disps0.1,
    prior.location=current.posterior.mean.prior.location.1,
    prior.scale=current.posterior.mean.prior.scale.1
  ) -
  (log(current.posterior.means0.1) +
     (log(current.posterior.means0.1) -
        log(proposed.posterior.means0.1))^2 / (2*mean.proposal.scales0))

current.posterior.means0.1[replace] <- proposed.posterior.means0.1[replace]
accept.means0.1[replace] <- accept.means0.1[replace] + 1/chain.length

# Chain 2
proposed.posterior.means0.2 <- rlnorm(
  n=genes, meanlog=log(current.posterior.means0.2), sdlog=sqrt.mean.proposal.scales0
)

replace <- log(runif(genes)) <=
  mean.conditional.log.posterior(
    n=samples0, sample.means=sample.means0,
    means=proposed.posterior.means0.2,
    disps=current.posterior.disps0.2,
    prior.location=current.posterior.mean.prior.location.2,
    prior.scale=current.posterior.mean.prior.scale.2
  ) +
  (log(proposed.posterior.means0.2) +
     (log(proposed.posterior.means0.2) -
        log(current.posterior.means0.2))^2 / (2*mean.proposal.scales0)) -
  mean.conditional.log.posterior(
    n=samples0, sample.means=sample.means0,
    means=current.posterior.means0.2,
    disps=current.posterior.disps0.2,
    prior.location=current.posterior.mean.prior.location.2,
    prior.scale=current.posterior.mean.prior.scale.2
  ) -
  (log(current.posterior.means0.2) +
     (log(current.posterior.means0.2) -
        log(proposed.posterior.means0.2))^2 / (2*mean.proposal.scales0))
```

```r
current.posterior.means0.2[replace] <- proposed.posterior.means0.2[replace]
accept.means0.2[replace] <- accept.means0.2[replace] + 1/chain.length

# Chain 3
proposed.posterior.means0.3 <- rlnorm(
  n=genes, meanlog=log(current.posterior.means0.3), sdlog=sqrt.mean.proposal.scales0
)

replace <- log(runif(genes)) <=
  mean.conditional.log.posterior(
    n=samples0,sample.means=sample.means0,
    means=proposed.posterior.means0.3,
    disps=current.posterior.disps0.3,
    prior.location=current.posterior.mean.prior.location.3,
    prior.scale=current.posterior.mean.prior.scale.3
  ) +
  (log(proposed.posterior.means0.3) +
     (log(proposed.posterior.means0.3) -
        log(current.posterior.means0.3))^2 / (2*mean.proposal.scales0)) -
  mean.conditional.log.posterior(
    n=samples0, sample.means=sample.means0,
    means=current.posterior.means0.3,
    disps=current.posterior.disps0.3,
    prior.location=current.posterior.mean.prior.location.3,
    prior.scale=current.posterior.mean.prior.scale.3
  ) -
  (log(current.posterior.means0.3) +
     (log(current.posterior.means0.3) -
        log(proposed.posterior.means0.3))^2 / (2*mean.proposal.scales0))

current.posterior.means0.3[replace] <- proposed.posterior.means0.3[replace]
accept.means0.3[replace] <- accept.means0.3[replace] + 1/chain.length

# Metropolis updates for group 1 per-gene means ####
# Chain 1
proposed.posterior.means1.1 <- rlnorm(
  n=genes, meanlog=log(current.posterior.means1.1), sdlog=sqrt.mean.proposal.scales1
)

replace <- log(runif(genes)) <=
  mean.conditional.log.posterior(
    n=samples1, sample.means=sample.means1,
    means=proposed.posterior.means1.1,
    disps=current.posterior.disps1.1,
    prior.location=current.posterior.mean.prior.location.1,
    prior.scale=current.posterior.mean.prior.scale.1
  ) +
  (log(proposed.posterior.means1.1) +
     (log(proposed.posterior.means1.1) -
        log(current.posterior.means1.1))^2 / (2*mean.proposal.scales1)) -
  mean.conditional.log.posterior(
    n=samples1, sample.means=sample.means1,
    means=current.posterior.means1.1,
```

```
      disps=current.posterior.disps1.1,
      prior.location=current.posterior.mean.prior.location.1,
      prior.scale=current.posterior.mean.prior.scale.1
    ) -
    (log(current.posterior.means1.1) +
      (log(current.posterior.means1.1) -
        log(proposed.posterior.means1.1))^2 / (2*mean.proposal.scales1))

  current.posterior.means1.1[replace] <- proposed.posterior.means1.1[replace]
  accept.means1.1[replace] <- accept.means1.1[replace] + 1/chain.length

  # Chain 2
  proposed.posterior.means1.2 <- rlnorm(
    n=genes, meanlog=log(current.posterior.means1.2), sdlog=sqrt.mean.proposal.scales1
  )

  replace <- log(runif(genes)) <=
    mean.conditional.log.posterior(
      n=samples1, sample.means=sample.means1,
      means=proposed.posterior.means1.2,
      disps=current.posterior.disps1.2,
      prior.location=current.posterior.mean.prior.location.2,
      prior.scale=current.posterior.mean.prior.scale.2
    ) +
    (log(proposed.posterior.means1.2) +
      (log(proposed.posterior.means1.2) -
        log(current.posterior.means1.2))^2 / (2*mean.proposal.scales1)) -
    mean.conditional.log.posterior(
      n=samples1,
      sample.means=sample.means1,
      means=current.posterior.means1.2,
      disps=current.posterior.disps1.2,
      prior.location=current.posterior.mean.prior.location.2,
      prior.scale=current.posterior.mean.prior.scale.2
    ) -
    (log(current.posterior.means1.2) +
      (log(current.posterior.means1.2) -
        log(proposed.posterior.means1.2))^2 / (2*mean.proposal.scales1))

  current.posterior.means1.2[replace] <- proposed.posterior.means1.2[replace]
  accept.means1.2[replace] <- accept.means1.2[replace] + 1/chain.length

  # Chain 3
  proposed.posterior.means1.3 <- rlnorm(
    n=genes, meanlog=log(current.posterior.means1.3), sdlog=sqrt.mean.proposal.scales1
  )

  replace <- log(runif(genes)) <=
    mean.conditional.log.posterior(
      n=samples1, sample.means=sample.means1,
      means=proposed.posterior.means1.3,
      disps=current.posterior.disps1.3,
      prior.location=current.posterior.mean.prior.location.3,
```

```
    prior.scale=current.posterior.mean.prior.scale.3
  ) +
  (log(proposed.posterior.means1.3) +
     (log(proposed.posterior.means1.3) -
        log(current.posterior.means1.3))^2 / (2*mean.proposal.scales1)) -
  mean.conditional.log.posterior(
    n=samples1, sample.means=sample.means1,
    means=current.posterior.means1.3,
    disps=current.posterior.disps1.3,
    prior.location=current.posterior.mean.prior.location.3,
    prior.scale=current.posterior.mean.prior.scale.3
  ) -
  (log(current.posterior.means1.3) +
     (log(current.posterior.means1.3) -
        log(proposed.posterior.means1.3))^2 / (2*mean.proposal.scales1))

current.posterior.means1.3[replace] <- proposed.posterior.means1.3[replace]
accept.means1.3[replace] <- accept.means1.3[replace] + 1/chain.length

# Metropolis updates for group 2 per-gene means ####
# Chain 1
proposed.posterior.means2.1 <- rlnorm(
  n=genes, meanlog=log(current.posterior.means2.1), sdlog=sqrt.mean.proposal.scales2
)

replace <- log(runif(genes)) <=
  mean.conditional.log.posterior(
    n=samples2, sample.means=sample.means2,
    means=proposed.posterior.means2.1,
    disps=current.posterior.disps2.1,
    prior.location=current.posterior.mean.prior.location.1,
    prior.scale=current.posterior.mean.prior.scale.1
  ) +
  (log(proposed.posterior.means2.1) +
     (log(proposed.posterior.means2.1) -
        log(current.posterior.means2.1))^2 / (2*mean.proposal.scales2)) -
  mean.conditional.log.posterior(
    n=samples2, sample.means=sample.means2,
    means=current.posterior.means2.1,
    disps=current.posterior.disps2.1,
    prior.location=current.posterior.mean.prior.location.1,
    prior.scale=current.posterior.mean.prior.scale.1
  ) -
  (log(current.posterior.means2.1) +
     (log(current.posterior.means2.1) -
        log(proposed.posterior.means2.1))^2 / (2*mean.proposal.scales2))

current.posterior.means2.1[replace] <- proposed.posterior.means2.1[replace]
accept.means2.1[replace] <- accept.means2.1[replace] + 1/chain.length

# Chain 2
proposed.posterior.means2.2 <- rlnorm(
  n=genes, meanlog=log(current.posterior.means2.2), sdlog=sqrt.mean.proposal.scales2
```

```
)
replace <- log(runif(genes)) <=
  mean.conditional.log.posterior(
    n=samples2, sample.means=sample.means2,
    means=proposed.posterior.means2.2,
    disps=current.posterior.disps2.2,
    prior.location=current.posterior.mean.prior.location.2,
    prior.scale=current.posterior.mean.prior.scale.2
  ) +
  (log(proposed.posterior.means2.2) +
     (log(proposed.posterior.means2.2) -
        log(current.posterior.means2.2))^2 / (2*mean.proposal.scales2)) -
  mean.conditional.log.posterior(
    n=samples2, sample.means=sample.means2,
    means=current.posterior.means2.2,
    disps=current.posterior.disps2.2,
    prior.location=current.posterior.mean.prior.location.2,
    prior.scale=current.posterior.mean.prior.scale.2
  ) -
  (log(current.posterior.means2.2) +
     (log(current.posterior.means2.2) -
        log(proposed.posterior.means2.2))^2 / (2*mean.proposal.scales2))

current.posterior.means2.2[replace] <-
  proposed.posterior.means2.2[replace]
accept.means2.2[replace] <-
  accept.means2.2[replace] + 1/chain.length

# Chain 3
proposed.posterior.means2.3 <- rlnorm(
  n=genes, meanlog=log(current.posterior.means2.3), sdlog=sqrt.mean.proposal.scales2
)

replace <- log(runif(genes)) <=
  mean.conditional.log.posterior(
    n=samples2, sample.means=sample.means2,
    means=proposed.posterior.means2.3,
    disps=current.posterior.disps2.3,
    prior.location=current.posterior.mean.prior.location.3,
    prior.scale=current.posterior.mean.prior.scale.3
  ) +
  (log(proposed.posterior.means2.3) +
     (log(proposed.posterior.means2.3) -
        log(current.posterior.means2.3))^2 / (2*mean.proposal.scales2)) -
  mean.conditional.log.posterior(
    n=samples2, sample.means=sample.means2,
    means=current.posterior.means2.3,
    disps=current.posterior.disps2.3,
    prior.location=current.posterior.mean.prior.location.3,
    prior.scale=current.posterior.mean.prior.scale.3
  ) -
  (log(current.posterior.means2.3) +
     (log(current.posterior.means2.3) -
```

```
        log(proposed.posterior.means2.3))^2 / (2*mean.proposal.scales2))

current.posterior.means2.3[replace] <- proposed.posterior.means2.3[replace]
accept.means2.3[replace] <- accept.means2.3[replace] + 1/chain.length

# Metropolis updates for overall per-gene dispersions ####
# Chain 1
proposed.posterior.disps0.1 <- rlnorm(
  n=genes, meanlog=log(current.posterior.disps0.1), sdlog=sqrt.disp.proposal.scales0
)

replace <- log(runif(genes)) <=
  disp.conditional.log.posterior(
    genes=genes, counts=counts, n=samples0,
    sample.means=sample.means0, means=current.posterior.means0.1,
    disps=proposed.posterior.disps0.1,
    prior.location=current.posterior.disp.prior.location.1,
    prior.scale=current.posterior.disp.prior.scale.1
  ) +
  (log(proposed.posterior.disps0.1) +
     (log(proposed.posterior.disps0.1) -
        log(current.posterior.disps0.1))^2 / (2*disp.proposal.scales0)) -
  disp.conditional.log.posterior(
    genes=genes, counts=counts, n=samples0,
    sample.means=sample.means0, means=current.posterior.means0.1,
    disps=current.posterior.disps0.1,
    prior.location=current.posterior.disp.prior.location.1,
    prior.scale=current.posterior.disp.prior.scale.1
  ) -
  (log(current.posterior.disps0.1) +
     (log(current.posterior.disps0.1) -
        log(proposed.posterior.disps0.1))^2 / (2*disp.proposal.scales0))

current.posterior.disps0.1[replace] <- proposed.posterior.disps0.1[replace]
accept.disps0.1[replace] <- accept.disps0.1[replace] + 1/chain.length

# Chain 2
proposed.posterior.disps0.2 <- rlnorm(
  n=genes, meanlog=log(current.posterior.disps0.2), sdlog=sqrt.disp.proposal.scales0
)

replace <- log(runif(genes)) <=
  disp.conditional.log.posterior(
    genes=genes, counts=counts, n=samples0,
    sample.means=sample.means0, means=current.posterior.means0.2,
    disps=proposed.posterior.disps0.2,
    prior.location=current.posterior.disp.prior.location.2,
    prior.scale=current.posterior.disp.prior.scale.2
  ) +
  (log(proposed.posterior.disps0.2) +
     (log(proposed.posterior.disps0.2) -
        log(current.posterior.disps0.2))^2 / (2*disp.proposal.scales0)) -
  disp.conditional.log.posterior(
```

```
      genes=genes, counts=counts, n=samples0,
      sample.means=sample.means0, means=current.posterior.means0.2,
      disps=current.posterior.disps0.2,
      prior.location=current.posterior.disp.prior.location.2,
      prior.scale=current.posterior.disp.prior.scale.2
    ) -
    (log(current.posterior.disps0.2) +
       (log(current.posterior.disps0.2) -
          log(proposed.posterior.disps0.2))^2 / (2*disp.proposal.scales0))

  current.posterior.disps0.2[replace] <- proposed.posterior.disps0.2[replace]
  accept.disps0.2[replace] <- accept.disps0.2[replace] + 1/chain.length

  # Chain 3
  proposed.posterior.disps0.3 <- rlnorm(
    n=genes, meanlog=log(current.posterior.disps0.3), sdlog=sqrt.disp.proposal.scales0
  )

  replace <- log(runif(genes)) <=
    disp.conditional.log.posterior(
      genes=genes, counts=counts, n=samples0,
      sample.means=sample.means0, means=current.posterior.means0.3,
      disps=proposed.posterior.disps0.3,
      prior.location=current.posterior.disp.prior.location.3,
      prior.scale=current.posterior.disp.prior.scale.3
    ) +
    (log(proposed.posterior.disps0.3) +
       (log(proposed.posterior.disps0.3) -
          log(current.posterior.disps0.3))^2 / (2*disp.proposal.scales0)) -
    disp.conditional.log.posterior(
      genes=genes, counts=counts, n=samples0,
      sample.means=sample.means0, means=current.posterior.means0.3,
      disps=current.posterior.disps0.3,
      prior.location=current.posterior.disp.prior.location.3,
      prior.scale=current.posterior.disp.prior.scale.3
    ) -
    (log(current.posterior.disps0.3) +
       (log(current.posterior.disps0.3) -
          log(proposed.posterior.disps0.3))^2 / (2*disp.proposal.scales0))

  current.posterior.disps0.3[replace] <- proposed.posterior.disps0.3[replace]
  accept.disps0.3[replace] <- accept.disps0.3[replace] + 1/chain.length

  # Metropolis updates for group 1 per-gene dispersions ####
  # Chain 1
  proposed.posterior.disps1.1 <- rlnorm(
    n=genes, meanlog=log(current.posterior.disps1.1), sdlog=sqrt.disp.proposal.scales1
  )

  replace <- log(runif(genes)) <=
    disp.conditional.log.posterior(
      genes=genes, counts=counts1, n=samples1,
      sample.means=sample.means1, means=current.posterior.means1.1,
```

```
      disps=proposed.posterior.disps1.1,
      prior.location=current.posterior.disp.prior.location.1,
      prior.scale=current.posterior.disp.prior.scale.1
    ) +
    (log(proposed.posterior.disps1.1) +
       (log(proposed.posterior.disps1.1) -
          log(current.posterior.disps1.1))^2 / (2*disp.proposal.scales1)) -
    disp.conditional.log.posterior(
      genes=genes, counts=counts1, n=samples1,
      sample.means=sample.means1, means=current.posterior.means1.1,
      disps=current.posterior.disps1.1,
      prior.location=current.posterior.disp.prior.location.1,
      prior.scale=current.posterior.disp.prior.scale.1
    ) -
    (log(current.posterior.disps1.1) +
       (log(current.posterior.disps1.1) -
          log(proposed.posterior.disps1.1))^2 / (2*disp.proposal.scales1))

  current.posterior.disps1.1[replace] <- proposed.posterior.disps1.1[replace]
  accept.disps1.1[replace] <- accept.disps1.1[replace] + 1/chain.length

  # Chain 2
  proposed.posterior.disps1.2 <- rlnorm(
    n=genes, meanlog=log(current.posterior.disps1.2), sdlog=sqrt.disp.proposal.scales1
  )
  replace <- log(runif(genes)) <=
    disp.conditional.log.posterior(
      genes=genes, counts=counts1, n=samples1,
      sample.means=sample.means1, means=current.posterior.means1.2,
      disps=proposed.posterior.disps1.2,
      prior.location=current.posterior.disp.prior.location.2,
      prior.scale=current.posterior.disp.prior.scale.2
    ) +
    (log(proposed.posterior.disps1.2) +
       (log(proposed.posterior.disps1.2) -
          log(current.posterior.disps1.2))^2 / (2*disp.proposal.scales1)) -
    disp.conditional.log.posterior(
      genes=genes, counts=counts1, n=samples1,
      sample.means=sample.means1, means=current.posterior.means1.2,
      disps=current.posterior.disps1.2,
      prior.location=current.posterior.disp.prior.location.2,
      prior.scale=current.posterior.disp.prior.scale.2
    ) -
    (log(current.posterior.disps1.2) +
       (log(current.posterior.disps1.2) -
          log(proposed.posterior.disps1.2))^2 / (2*disp.proposal.scales1))

  current.posterior.disps1.2[replace] <- proposed.posterior.disps1.2[replace]
  accept.disps1.2[replace] <- accept.disps1.2[replace] + 1/chain.length

  # Chain 3
  proposed.posterior.disps1.3 <- rlnorm(
    n=genes, meanlog=log(current.posterior.disps1.3), sdlog=sqrt.disp.proposal.scales1
```

```
  )

  replace <- log(runif(genes)) <=
    disp.conditional.log.posterior(
      genes=genes, counts=counts1, n=samples1,
      sample.means=sample.means1, means=current.posterior.means1.3,
      disps=proposed.posterior.disps1.3,
      prior.location=current.posterior.disp.prior.location.3,
      prior.scale=current.posterior.disp.prior.scale.3
    ) +
    (log(proposed.posterior.disps1.3) +
       (log(proposed.posterior.disps1.3) -
          log(current.posterior.disps1.3))^2 / (2*disp.proposal.scales1)) -
    disp.conditional.log.posterior(
      genes=genes, counts=counts1, n=samples1,
      sample.means=sample.means1, means=current.posterior.means1.3,
      disps=current.posterior.disps1.3,
      prior.location=current.posterior.disp.prior.location.3,
      prior.scale=current.posterior.disp.prior.scale.3
    ) -
    (log(current.posterior.disps1.3) +
       (log(current.posterior.disps1.3) -
          log(proposed.posterior.disps1.3))^2 / (2*disp.proposal.scales1))

  current.posterior.disps1.3[replace] <- proposed.posterior.disps1.3[replace]
  accept.disps1.3[replace] <- accept.disps1.3[replace] + 1/chain.length

  # Metropolis updates for group 2 per-gene dispersions ####
  # Chain 1
  proposed.posterior.disps2.1 <- rlnorm(
    n=genes, meanlog=log(current.posterior.disps2.1), sdlog=sqrt.disp.proposal.scales2
  )
  replace <- log(runif(genes)) <=
    disp.conditional.log.posterior(
      genes=genes, counts=counts2, n=samples2,
      sample.means=sample.means2, means=current.posterior.means2.1,
      disps=proposed.posterior.disps2.1,
      prior.location=current.posterior.disp.prior.location.1,
      prior.scale=current.posterior.disp.prior.scale.1
    ) +
    (log(proposed.posterior.disps2.1) +
       (log(proposed.posterior.disps2.1) -
          log(current.posterior.disps2.1))^2 / (2*disp.proposal.scales2)) -
    disp.conditional.log.posterior(
      genes=genes, counts=counts2, n=samples2,
      sample.means=sample.means2, means=current.posterior.means2.1,
      disps=current.posterior.disps2.1,
      prior.location=current.posterior.disp.prior.location.1,
      prior.scale=current.posterior.disp.prior.scale.1
    ) -
    (log(current.posterior.disps2.1) +
       (log(current.posterior.disps2.1) -
          log(proposed.posterior.disps2.1))^2 / (2*disp.proposal.scales2))
```

```
current.posterior.disps2.1[replace] <- proposed.posterior.disps2.1[replace]
accept.disps2.1[replace] <- accept.disps2.1[replace] + 1/chain.length

# Chain 2
proposed.posterior.disps2.2 <- rlnorm(
  n=genes, meanlog=log(current.posterior.disps2.2),
  sdlog=sqrt.disp.proposal.scales2
)
replace <- log(runif(genes)) <=
  disp.conditional.log.posterior(
    genes=genes, counts=counts2, n=samples2,
    sample.means=sample.means2, means=current.posterior.means2.2,
    disps=proposed.posterior.disps2.2,
    prior.location=current.posterior.disp.prior.location.2,
    prior.scale=current.posterior.disp.prior.scale.2
  ) +
  (log(proposed.posterior.disps2.2) +
     (log(proposed.posterior.disps2.2) -
        log(current.posterior.disps2.2))^2 / (2*disp.proposal.scales2)) -
  disp.conditional.log.posterior(
    genes=genes, counts=counts2,n=samples2,
    sample.means=sample.means2, means=current.posterior.means2.2,
    disps=current.posterior.disps2.2,
    prior.location=current.posterior.disp.prior.location.2,
    prior.scale=current.posterior.disp.prior.scale.2
  ) -
  (log(current.posterior.disps2.2) +
     (log(current.posterior.disps2.2) -
        log(proposed.posterior.disps2.2))^2 / (2*disp.proposal.scales2))

current.posterior.disps2.2[replace] <- proposed.posterior.disps2.2[replace]
accept.disps2.2[replace] <- accept.disps2.2[replace] + 1/chain.length

# Chain 3
proposed.posterior.disps2.3 <- rlnorm(
  n=genes, meanlog=log(current.posterior.disps2.3), sdlog=sqrt.disp.proposal.scales2
)

replace <- log(runif(genes)) <=
  disp.conditional.log.posterior(
    genes=genes, counts=counts2, n=samples2,
    sample.means=sample.means2, means=current.posterior.means2.3,
    disps=proposed.posterior.disps2.3,
    prior.location=current.posterior.disp.prior.location.3,
    prior.scale=current.posterior.disp.prior.scale.3
  ) +
  (log(proposed.posterior.disps2.3) +
     (log(proposed.posterior.disps2.3) -
        log(current.posterior.disps2.3))^2 / (2*disp.proposal.scales2)) -
  disp.conditional.log.posterior(
    genes=genes, counts=counts2, n=samples2,
    sample.means=sample.means2, means=current.posterior.means2.3,
    disps=current.posterior.disps2.3,
```

```
      prior.location=current.posterior.disp.prior.location.3,
      prior.scale=current.posterior.disp.prior.scale.3
    ) -
    (log(current.posterior.disps2.3) +
      (log(current.posterior.disps2.3) -
        log(proposed.posterior.disps2.3))^2 / (2*disp.proposal.scales2))

  current.posterior.disps2.3[replace] <- proposed.posterior.disps2.3[replace]
  accept.disps2.3[replace] <- accept.disps2.3[replace] + 1/chain.length

  # Gibbs update for prior location parameter for mean ####
  current.posterior.mean.prior.location.1 <- rnorm(
    n=1,
    mean=prior.location.posterior.mean(
      hyperprior.mean=2.5, hyperprior.var=20,
      prior.scale=current.posterior.mean.prior.scale.1,
      parameters0=current.posterior.means0.1,
      parameters1=current.posterior.means1.1,
      parameters2=current.posterior.means2.1,
      z=current.posterior.indicators.1
    ),
    sd=prior.location.posterior.sd(
      hyperprior.var=20,
      prior.scale=current.posterior.mean.prior.scale.1,
      z=current.posterior.indicators.1
    )
  )

  current.posterior.mean.prior.location.2 <- rnorm(
    n=1,
    mean=prior.location.posterior.mean(
      hyperprior.mean=2.5, hyperprior.var=20,
      prior.scale=current.posterior.mean.prior.scale.2,
      parameters0=current.posterior.means0.2,
      parameters1=current.posterior.means1.2,
      parameters2=current.posterior.means2.2,
      z=current.posterior.indicators.2
    ),
    sd=prior.location.posterior.sd(
      hyperprior.var=20,
      prior.scale=current.posterior.mean.prior.scale.2,
      z=current.posterior.indicators.2
    )
  )

  current.posterior.mean.prior.location.3 <- rnorm(
    n=1,
    mean=prior.location.posterior.mean(
      hyperprior.mean=2.5, hyperprior.var=20,
      prior.scale=current.posterior.mean.prior.scale.3,
      parameters0=current.posterior.means0.3,
      parameters1=current.posterior.means1.3,
      parameters2=current.posterior.means2.3,
```

```
      z=current.posterior.indicators.3
    ),
    sd=prior.location.posterior.sd(
      hyperprior.var=20,
      prior.scale=current.posterior.mean.prior.scale.3,
      z=current.posterior.indicators.3
    )
  )

  # Metropolis update for prior scale parameter for mean ####
  # Chain 1
  proposed.posterior.mean.prior.scale.1 <- rnorm(
    n=1, mean=current.posterior.mean.prior.scale.1, sd=mean.prior.scale.proposal.sd
  )
  if (log(runif(1)) <=
      mean.prior.scale.log.posterior(
        prior.scale=proposed.posterior.mean.prior.scale.1,
        prior.location=current.posterior.mean.prior.location.1,
        g=genes, means0=current.posterior.means0.1,
        means1=current.posterior.means1.1,
        means2=current.posterior.means2.1,
        z=current.posterior.indicators.1
      ) -
      mean.prior.scale.log.posterior(
        prior.scale=current.posterior.mean.prior.scale.1,
        prior.location=current.posterior.mean.prior.location.1,
        g=genes, means0=current.posterior.means0.1,
        means1=current.posterior.means1.1,
        means2=current.posterior.means2.1,
        z=current.posterior.indicators.1
      )) {
    current.posterior.mean.prior.scale.1 <- proposed.posterior.mean.prior.scale.1
    accept.mean.prior.scale.1 <- accept.mean.prior.scale.1 + 1/chain.length
  }

  # Chain 2
  proposed.posterior.mean.prior.scale.2 <- rnorm(
    n=1, mean=current.posterior.mean.prior.scale.2, sd=mean.prior.scale.proposal.sd
  )
  if (log(runif(1)) <=
      mean.prior.scale.log.posterior(
        prior.scale=proposed.posterior.mean.prior.scale.2,
        prior.location=current.posterior.mean.prior.location.2,
        g=genes, means0=current.posterior.means0.2,
        means1=current.posterior.means1.2,
        means2=current.posterior.means2.2,
        z=current.posterior.indicators.2
      ) -
      mean.prior.scale.log.posterior(
        prior.scale=current.posterior.mean.prior.scale.2,
        prior.location=current.posterior.mean.prior.location.2,
        g=genes, means0=current.posterior.means0.2,
        means1=current.posterior.means1.2,
```

```
      means2=current.posterior.means2.2,
      z=current.posterior.indicators.2
    )) {
  current.posterior.mean.prior.scale.2 <- proposed.posterior.mean.prior.scale.2
  accept.mean.prior.scale.2 <- accept.mean.prior.scale.2 + 1/chain.length
}

# Chain 3
proposed.posterior.mean.prior.scale.3 <- rnorm(
  n=1, mean=current.posterior.mean.prior.scale.3, sd=mean.prior.scale.proposal.sd
)
if (log(runif(1)) <=
    mean.prior.scale.log.posterior(
      prior.scale=proposed.posterior.mean.prior.scale.3,
      prior.location=current.posterior.mean.prior.location.3,
      g=genes, means0=current.posterior.means0.3,
      means1=current.posterior.means1.3,
      means2=current.posterior.means2.3,
      z=current.posterior.indicators.3
    ) -
    mean.prior.scale.log.posterior(
      prior.scale=current.posterior.mean.prior.scale.3,
      prior.location=current.posterior.mean.prior.location.3,
      g=genes, means0=current.posterior.means0.3,
      means1=current.posterior.means1.3,
      means2=current.posterior.means2.3,
      z=current.posterior.indicators.3
    )) {
  current.posterior.mean.prior.scale.3 <- proposed.posterior.mean.prior.scale.3
  accept.mean.prior.scale.3 <- accept.mean.prior.scale.3 + 1/chain.length
}

# Gibbs update for prior location parameter for dispersion ####
current.posterior.disp.prior.location.1 <- rnorm(
  n=1,
  mean=prior.location.posterior.mean(
    hyperprior.mean=-2.5, hyperprior.var=2,
    prior.scale=current.posterior.disp.prior.scale.1,
    parameters0=current.posterior.disps0.1,
    parameters1=current.posterior.disps1.1,
    parameters2=current.posterior.disps2.1,
    z=current.posterior.indicators.1
  ),
  sd=prior.location.posterior.sd(
    hyperprior.var=2,
    prior.scale=current.posterior.disp.prior.scale.1,
    z=current.posterior.indicators.1
  )
)

current.posterior.disp.prior.location.2 <- rnorm(
  n=1,
  mean=prior.location.posterior.mean(
```

```r
      hyperprior.mean=-2.5, hyperprior.var=2,
      prior.scale=current.posterior.disp.prior.scale.2,
      parameters0=current.posterior.disps0.2,
      parameters1=current.posterior.disps1.2,
      parameters2=current.posterior.disps2.2,
      z=current.posterior.indicators.2
    ),
    sd=prior.location.posterior.sd(
      hyperprior.var=2,
      prior.scale=current.posterior.disp.prior.scale.2,
      z=current.posterior.indicators.2
    )
  )

  current.posterior.disp.prior.location.3 <- rnorm(
    n=1,
    mean=prior.location.posterior.mean(
      hyperprior.mean=-2.5, hyperprior.var=2,
      prior.scale=current.posterior.disp.prior.scale.3,
      parameters0=current.posterior.disps0.3,
      parameters1=current.posterior.disps1.3,
      parameters2=current.posterior.disps2.3,
      z=current.posterior.indicators.3
    ),
    sd=prior.location.posterior.sd(
      hyperprior.var=2,
      prior.scale=current.posterior.disp.prior.scale.3,
      z=current.posterior.indicators.3
    )
  )

  # Metropolis update for prior scale parameter for dispersion ####
  # Chain 1
  proposed.posterior.disp.prior.scale.1 <- rnorm(
    n=1, mean=current.posterior.disp.prior.scale.1, sd=disp.prior.scale.proposal.sd
  )
  if (log(runif(1)) <=
      disp.prior.scale.log.posterior(
        prior.scale=proposed.posterior.disp.prior.scale.1,
        prior.location=current.posterior.disp.prior.location.1,
        g=genes, disps0=current.posterior.disps0.1,
        disps1=current.posterior.disps1.1,
        disps2=current.posterior.disps2.1,
        z=current.posterior.indicators.1
      ) -
      disp.prior.scale.log.posterior(
        prior.scale=current.posterior.disp.prior.scale.1,
        prior.location=current.posterior.disp.prior.location.1,
        g=genes, disps0=current.posterior.disps0.1,
        disps1=current.posterior.disps1.1,
        disps2=current.posterior.disps2.1,
        z=current.posterior.indicators.1
      )) {
```

```
    current.posterior.disp.prior.scale.1 <- proposed.posterior.disp.prior.scale.1
    accept.disp.prior.scale.1 <- accept.disp.prior.scale.1 + 1/chain.length
  }

  # Chain 2
  proposed.posterior.disp.prior.scale.2 <- rnorm(
    n=1, mean=current.posterior.disp.prior.scale.2, sd=disp.prior.scale.proposal.sd
  )
  if (log(runif(1)) <=
      disp.prior.scale.log.posterior(
        prior.scale=proposed.posterior.disp.prior.scale.2,
        prior.location=current.posterior.disp.prior.location.2,
        g=genes, disps0=current.posterior.disps0.2,
        disps1=current.posterior.disps1.2,
        disps2=current.posterior.disps2.2,
        z=current.posterior.indicators.2
      ) -
      disp.prior.scale.log.posterior(
        prior.scale=current.posterior.disp.prior.scale.2,
        prior.location=current.posterior.disp.prior.location.2,
        g=genes, disps0=current.posterior.disps0.2,
        disps1=current.posterior.disps1.2,
        disps2=current.posterior.disps2.2,
        z=current.posterior.indicators.2
      )) {
    current.posterior.disp.prior.scale.2 <- proposed.posterior.disp.prior.scale.2
    accept.disp.prior.scale.2 <- accept.disp.prior.scale.2 + 1/chain.length
  }

  # Chain 3
  proposed.posterior.disp.prior.scale.3 <- rnorm(
    n=1, mean=current.posterior.disp.prior.scale.3, sd=disp.prior.scale.proposal.sd
  )
  if (log(runif(1)) <=
      disp.prior.scale.log.posterior(
        prior.scale=proposed.posterior.disp.prior.scale.3,
        prior.location=current.posterior.disp.prior.location.3,
        g=genes, disps0=current.posterior.disps0.3,
        disps1=current.posterior.disps1.3,
        disps2=current.posterior.disps2.3,
        z=current.posterior.indicators.3
      ) -
      disp.prior.scale.log.posterior(
        prior.scale=current.posterior.disp.prior.scale.3,
        prior.location=current.posterior.disp.prior.location.3,
        g=genes, disps0=current.posterior.disps0.3,
        disps1=current.posterior.disps1.3,
        disps2=current.posterior.disps2.3,
        z=current.posterior.indicators.3
      )) {
    current.posterior.disp.prior.scale.3 <- proposed.posterior.disp.prior.scale.3
    accept.disp.prior.scale.3 <- accept.disp.prior.scale.3 + 1/chain.length
  }
```

```
# Gibbs updates for per-gene mixture components ####
current.posterior.indicators.1 <- rbinom(
  n=genes, size=1,
  prob=posterior.indicator.probabilities(
    genes=genes, counts=counts, counts1=counts1, counts2=counts2,
    n=samples0, n1=samples1, n2=samples2, sample.means0=sample.means0,
    sample.means1=sample.means1, sample.means2=sample.means2,
    means0=current.posterior.means0.1, means1=current.posterior.means1.1,
    means2=current.posterior.means2.1, disps0=current.posterior.disps0.1,
    disps1=current.posterior.disps1.1, disps2=current.posterior.disps2.1,
    mean.prior.location=current.posterior.mean.prior.location.1,
    mean.prior.scale=current.posterior.mean.prior.scale.1,
    disp.prior.location=current.posterior.disp.prior.location.1,
    disp.prior.scale=current.posterior.disp.prior.scale.1,
    lambda=current.posterior.proportion.1
  )
)

current.posterior.indicators.2 <- rbinom(
  n=genes, size=1,
  prob=posterior.indicator.probabilities(
    genes=genes, counts=counts, counts1=counts1, counts2=counts2,
    n=samples0, n1=samples1, n2=samples2, sample.means0=sample.means0,
    sample.means1=sample.means1, sample.means2=sample.means2,
    means0=current.posterior.means0.2, means1=current.posterior.means1.2,
    means2=current.posterior.means2.2, disps0=current.posterior.disps0.2,
    disps1=current.posterior.disps1.2, disps2=current.posterior.disps2.2,
    mean.prior.location=current.posterior.mean.prior.location.2,
    mean.prior.scale=current.posterior.mean.prior.scale.2,
    disp.prior.location=current.posterior.disp.prior.location.2,
    disp.prior.scale=current.posterior.disp.prior.scale.2,
    lambda=current.posterior.proportion.2
  )
)

current.posterior.indicators.3 <- rbinom(
  n=genes, size=1,
  prob=posterior.indicator.probabilities(
    genes=genes, counts=counts, counts1=counts1, counts2=counts2,
    n=samples0, n1=samples1, n2=samples2, sample.means0=sample.means0,
    sample.means1=sample.means1, sample.means2=sample.means2,
    means0=current.posterior.means0.3, means1=current.posterior.means1.3,
    means2=current.posterior.means2.3, disps0=current.posterior.disps0.3,
    disps1=current.posterior.disps1.3, disps2=current.posterior.disps2.3,
    mean.prior.location=current.posterior.mean.prior.location.3,
    mean.prior.scale=current.posterior.mean.prior.scale.3,
    disp.prior.location=current.posterior.disp.prior.location.3,
    disp.prior.scale=current.posterior.disp.prior.scale.3,
    lambda=current.posterior.proportion.3
  )
)

# Gibbs update for mixture proportion ####
```

```
current.posterior.proportion.1 <- rbeta(
  n=1, shape1=1 + sum(current.posterior.indicators.1),
  shape2=1 + genes - sum(current.posterior.indicators.1)
)

current.posterior.proportion.2 <- rbeta(
  n=1, shape1=1 + sum(current.posterior.indicators.2),
  shape2=1 + genes - sum(current.posterior.indicators.2)
)

current.posterior.proportion.3 <- rbeta(
  n=1, shape1=1 + sum(current.posterior.indicators.3),
  shape2=1 + genes - sum(current.posterior.indicators.3)
)

# Update posterior samples ####
if (iter/thin==round(iter/thin)) {
  posterior.means0.1[iter/thin,] <- current.posterior.means0.1
  posterior.means1.1[iter/thin,] <- current.posterior.means1.1
  posterior.means2.1[iter/thin,] <- current.posterior.means2.1
  posterior.means0.2[iter/thin,] <- current.posterior.means0.2
  posterior.means1.2[iter/thin,] <- current.posterior.means1.2
  posterior.means2.2[iter/thin,] <- current.posterior.means2.2
  posterior.means0.3[iter/thin,] <- current.posterior.means0.3
  posterior.means1.3[iter/thin,] <- current.posterior.means1.3
  posterior.means2.3[iter/thin,] <- current.posterior.means2.3
  posterior.disps0.1[iter/thin,] <- current.posterior.disps0.1
  posterior.disps1.1[iter/thin,] <- current.posterior.disps1.1
  posterior.disps2.1[iter/thin,] <- current.posterior.disps2.1
  posterior.disps0.2[iter/thin,] <- current.posterior.disps0.2
  posterior.disps1.2[iter/thin,] <- current.posterior.disps1.2
  posterior.disps2.2[iter/thin,] <- current.posterior.disps2.2
  posterior.disps0.3[iter/thin,] <- current.posterior.disps0.3
  posterior.disps1.3[iter/thin,] <- current.posterior.disps1.3
  posterior.disps2.3[iter/thin,] <- current.posterior.disps2.3
  posterior.mean.prior.location.1[iter/thin] <-
    current.posterior.mean.prior.location.1
  posterior.mean.prior.location.2[iter/thin] <-
    current.posterior.mean.prior.location.2
  posterior.mean.prior.location.3[iter/thin] <-
    current.posterior.mean.prior.location.3
  posterior.mean.prior.scale.1[iter/thin] <- current.posterior.mean.prior.scale.1
  posterior.mean.prior.scale.2[iter/thin] <- current.posterior.mean.prior.scale.2
  posterior.mean.prior.scale.3[iter/thin] <- current.posterior.mean.prior.scale.3
  posterior.disp.prior.location.1[iter/thin] <-
    current.posterior.disp.prior.location.1
  posterior.disp.prior.location.2[iter/thin] <-
    current.posterior.disp.prior.location.2
  posterior.disp.prior.location.3[iter/thin] <-
    current.posterior.disp.prior.location.3
  posterior.disp.prior.scale.1[iter/thin] <- current.posterior.disp.prior.scale.1
  posterior.disp.prior.scale.2[iter/thin] <- current.posterior.disp.prior.scale.2
  posterior.disp.prior.scale.3[iter/thin] <- current.posterior.disp.prior.scale.3
```

```
    posterior.indicators.1[iter/thin,] <- current.posterior.indicators.1
    posterior.indicators.2[iter/thin,] <- current.posterior.indicators.2
    posterior.indicators.3[iter/thin,] <- current.posterior.indicators.3
    posterior.proportion.1[iter/thin] <- current.posterior.proportion.1
    posterior.proportion.2[iter/thin] <- current.posterior.proportion.2
    posterior.proportion.3[iter/thin] <- current.posterior.proportion.3
  }

}

return(list(
  "chain.length"=chain.length, "thin"=thin,
  "inits1"=inits1, "inits2"=inits2, "inits3"=inits3,
  "mean.proposal.scales0"=mean.proposal.scales0,
  "mean.proposal.scales1"=mean.proposal.scales1,
  "mean.proposal.scales2"=mean.proposal.scales2,
  "disp.proposal.scales0"=disp.proposal.scales0,
  "disp.proposal.scales1"=disp.proposal.scales1,
  "disp.proposal.scales2"=disp.proposal.scales2,
  "mean.prior.scale.proposal.sd"=mean.prior.scale.proposal.sd,
  "disp.prior.scale.proposal.sd"=disp.prior.scale.proposal.sd,
  "accept.means0.1"=accept.means0.1,
  "accept.means1.1"=accept.means1.1,
  "accept.means2.1"=accept.means2.1,
  "accept.means0.2"=accept.means0.2,
  "accept.means1.2"=accept.means1.2,
  "accept.means2.2"=accept.means2.2,
  "accept.means0.3"=accept.means0.3,
  "accept.means1.3"=accept.means1.3,
  "accept.means2.3"=accept.means2.3,
  "accept.disps0.1"=accept.disps0.1,
  "accept.disps1.1"=accept.disps1.1,
  "accept.disps2.1"=accept.disps2.1,
  "accept.disps0.2"=accept.disps0.2,
  "accept.disps1.2"=accept.disps1.2,
  "accept.disps2.2"=accept.disps2.2,
  "accept.disps0.3"=accept.disps0.3,
  "accept.disps1.3"=accept.disps1.3,
  "accept.disps2.3"=accept.disps2.3,
  "accept.mean.prior.scale.1"=accept.mean.prior.scale.1,
  "accept.mean.prior.scale.2"=accept.mean.prior.scale.2,
  "accept.mean.prior.scale.3"=accept.mean.prior.scale.3,
  "accept.disp.prior.scale.1"=accept.disp.prior.scale.1,
  "accept.disp.prior.scale.2"=accept.disp.prior.scale.2,
  "accept.disp.prior.scale.3"=accept.disp.prior.scale.3,
  "posterior.means0.1"=posterior.means0.1,
  "posterior.means1.1"=posterior.means1.1,
  "posterior.means2.1"=posterior.means2.1,
  "posterior.means0.2"=posterior.means0.2,
  "posterior.means1.2"=posterior.means1.2,
  "posterior.means2.2"=posterior.means2.2,
  "posterior.means0.3"=posterior.means0.3,
  "posterior.means1.3"=posterior.means1.3,
```

```
    "posterior.means2.3"=posterior.means2.3,
    "posterior.disps0.1"=posterior.disps0.1,
    "posterior.disps1.1"=posterior.disps1.1,
    "posterior.disps2.1"=posterior.disps2.1,
    "posterior.disps0.2"=posterior.disps0.2,
    "posterior.disps1.2"=posterior.disps1.2,
    "posterior.disps2.2"=posterior.disps2.2,
    "posterior.disps0.3"=posterior.disps0.3,
    "posterior.disps1.3"=posterior.disps1.3,
    "posterior.disps2.3"=posterior.disps2.3,
    "posterior.mean.prior.location.1"=posterior.mean.prior.location.1,
    "posterior.mean.prior.location.2"=posterior.mean.prior.location.2,
    "posterior.mean.prior.location.3"=posterior.mean.prior.location.3,
    "posterior.mean.prior.scale.1"=posterior.mean.prior.scale.1,
    "posterior.mean.prior.scale.2"=posterior.mean.prior.scale.2,
    "posterior.mean.prior.scale.3"=posterior.mean.prior.scale.3,
    "posterior.disp.prior.location.1"=posterior.disp.prior.location.1,
    "posterior.disp.prior.location.2"=posterior.disp.prior.location.2,
    "posterior.disp.prior.location.3"=posterior.disp.prior.location.3,
    "posterior.disp.prior.scale.1"=posterior.disp.prior.scale.1,
    "posterior.disp.prior.scale.2"=posterior.disp.prior.scale.2,
    "posterior.disp.prior.scale.3"=posterior.disp.prior.scale.3,
    "posterior.indicators.1"=posterior.indicators.1,
    "posterior.indicators.2"=posterior.indicators.2,
    "posterior.indicators.3"=posterior.indicators.3,
    "posterior.proportion.1"=posterior.proportion.1,
    "posterior.proportion.2"=posterior.proportion.2,
    "posterior.proportion.3"=posterior.proportion.3
    ))

}
```

##### Function to run one MCMC chain

```
ln_hmm_1_chain <- function(
  counts, groups, chain.length, thin=1, inits,
  mean.proposal.scales0=rep(0.2, ncol(counts)),
  mean.proposal.scales1=rep(0.2, ncol(counts)),
  mean.proposal.scales2=rep(0.2, ncol(counts)),
  disp.proposal.scales0=rep(0.5, ncol(counts)),
  disp.proposal.scales1=rep(0.5, ncol(counts)),
  disp.proposal.scales2=rep(0.5, ncol(counts)),
  mean.prior.scale.proposal.sd=0.1, disp.prior.scale.proposal.sd=0.4
  ) {

  genes <- ncol(counts)
  counts1 <- counts[groups==1,]
  counts2 <- counts[groups==2,]
  samples0 <- nrow(counts)
  samples1 <- nrow(counts1)
  samples2 <- nrow(counts2)
  sample.means0 <- colMeans(counts)
```

```r
sample.means1 <- colMeans(counts1)
sample.means2 <- colMeans(counts2)
sqrt.mean.proposal.scales0 <- sqrt(mean.proposal.scales0)
sqrt.mean.proposal.scales1 <- sqrt(mean.proposal.scales1)
sqrt.mean.proposal.scales2 <- sqrt(mean.proposal.scales2)
sqrt.disp.proposal.scales0 <- sqrt(disp.proposal.scales0)
sqrt.disp.proposal.scales1 <- sqrt(disp.proposal.scales1)
sqrt.disp.proposal.scales2 <- sqrt(disp.proposal.scales2)

# Create empty posterior sample matrices and vectors ####
posterior.means0 <- matrix(nrow=chain.length/thin, ncol=genes)
posterior.means1 <- matrix(nrow=chain.length/thin, ncol=genes)
posterior.means2 <- matrix(nrow=chain.length/thin, ncol=genes)
posterior.disps0 <- matrix(nrow=chain.length/thin, ncol=genes)
posterior.disps1 <- matrix(nrow=chain.length/thin, ncol=genes)
posterior.disps2 <- matrix(nrow=chain.length/thin, ncol=genes)
posterior.mean.prior.location <- numeric(chain.length/thin)
posterior.mean.prior.scale <- numeric(chain.length/thin)
posterior.disp.prior.location <- numeric(chain.length/thin)
posterior.disp.prior.scale <- numeric(chain.length/thin)
posterior.indicators <- matrix(nrow=chain.length/thin, ncol=genes)
posterior.proportion <- numeric(chain.length/thin)

# Initial values ####
current.posterior.means0 <- inits$means0
current.posterior.means1 <- inits$means1
current.posterior.means2 <- inits$means2
current.posterior.disps0 <- inits$disps0
current.posterior.disps1 <- inits$disps1
current.posterior.disps2 <- inits$disps2
current.posterior.mean.prior.location <- inits$mean.prior.location
current.posterior.mean.prior.scale <- inits$mean.prior.scale
current.posterior.disp.prior.location <- inits$mean.prior.location
current.posterior.disp.prior.scale <- inits$disp.prior.scale
current.posterior.indicators <- numeric(genes)
current.posterior.proportion = 0.5

# Create acceptance rate vectors/variables ####
accept.means0 <- numeric(genes)
accept.means1 <- numeric(genes)
accept.means2 <- numeric(genes)
accept.disps0 <- numeric(genes)
accept.disps1 <- numeric(genes)
accept.disps2 <- numeric(genes)
accept.mean.prior.scale <- 0
accept.disp.prior.scale <- 0

# Run MCMC ####
for (iter in 1:chain.length) {
  # Metropolis updates for per-gene overall means ####
  proposed.posterior.means0 <- rlnorm(
    n=genes, meanlog=log(current.posterior.means0), sdlog=sqrt.mean.proposal.scales0
  )
```

```
replace <- log(runif(genes)) <=
  mean.conditional.log.posterior(
    n=samples0, sample.means=sample.means0,
    means=proposed.posterior.means0,
    disps=current.posterior.disps0,
    prior.location=current.posterior.mean.prior.location,
    prior.scale=current.posterior.mean.prior.scale
  ) +
  (log(proposed.posterior.means0) +
     (log(proposed.posterior.means0) -
        log(current.posterior.means0))^2 / (2*mean.proposal.scales0)) -
  mean.conditional.log.posterior(
    n=samples0,
    sample.means=sample.means0,
    means=current.posterior.means0,
    disps=current.posterior.disps0,
    prior.location=current.posterior.mean.prior.location,
    prior.scale=current.posterior.mean.prior.scale
  ) -
  (log(current.posterior.means0) +
     (log(current.posterior.means0) -
        log(proposed.posterior.means0))^2 / (2*mean.proposal.scales0))

current.posterior.means0[replace] <- proposed.posterior.means0[replace]
accept.means0[replace] <- accept.means0[replace] + 1/chain.length

# Metropolis updates for per-gene group 1 means ####
proposed.posterior.means1 <- rlnorm(
  n=genes, meanlog=log(current.posterior.means1), sdlog=sqrt.mean.proposal.scales1
)
replace <- log(runif(genes)) <=
  mean.conditional.log.posterior(
    n=samples1, sample.means=sample.means1,
    means=proposed.posterior.means1,
    disps=current.posterior.disps1,
    prior.location=current.posterior.mean.prior.location,
    prior.scale=current.posterior.mean.prior.scale
  ) +
  (log(proposed.posterior.means1) +
     (log(proposed.posterior.means1) -
        log(current.posterior.means1))^2 / (2*mean.proposal.scales1)) -
  mean.conditional.log.posterior(
    n=samples1, sample.means=sample.means1,
    means=current.posterior.means1,
    disps=current.posterior.disps1,
    prior.location=current.posterior.mean.prior.location,
    prior.scale=current.posterior.mean.prior.scale
  ) -
  (log(current.posterior.means1) +
     (log(current.posterior.means1) -
        log(proposed.posterior.means1))^2 / (2*mean.proposal.scales1))

current.posterior.means1[replace] <- proposed.posterior.means1[replace]
```

```
  accept.means1[replace] <- accept.means1[replace] + 1/chain.length

  # Metropolis updates for per-gene group 2 means ####
  proposed.posterior.means2 <- rlnorm(
    n=genes, meanlog=log(current.posterior.means2), sdlog=sqrt.mean.proposal.scales2
  )

  replace <- log(runif(genes)) <=
    mean.conditional.log.posterior(
      n=samples2, sample.means=sample.means2,
      means=proposed.posterior.means2,
      disps=current.posterior.disps2,
      prior.location=current.posterior.mean.prior.location,
      prior.scale=current.posterior.mean.prior.scale
    ) +
    (log(proposed.posterior.means2) +
       (log(proposed.posterior.means2) -
          log(current.posterior.means2))^2 / (2*mean.proposal.scales2)) -
    mean.conditional.log.posterior(
      n=samples2,
      sample.means=sample.means2,
      means=current.posterior.means2,
      disps=current.posterior.disps2,
      prior.location=current.posterior.mean.prior.location,
      prior.scale=current.posterior.mean.prior.scale
    ) -
    (log(current.posterior.means2) +
       (log(current.posterior.means2) -
          log(proposed.posterior.means2))^2 / (2*mean.proposal.scales2))

  current.posterior.means2[replace] <- proposed.posterior.means2[replace]
  accept.means2[replace] <- accept.means2[replace] + 1/chain.length

  # Metropolis updates for per-gene overall dispersions ####
  proposed.posterior.disps0 <- rlnorm(
    n=genes, meanlog=log(current.posterior.disps0), sdlog=sqrt.disp.proposal.scales0
  )

  replace <- log(runif(genes)) <=
    disp.conditional.log.posterior(
      genes=genes, counts=counts, n=samples0,
      sample.means=sample.means0, means=current.posterior.means0,
      disps=proposed.posterior.disps0,
      prior.location=current.posterior.disp.prior.location,
      prior.scale=current.posterior.disp.prior.scale
    ) +
    (log(proposed.posterior.disps0) +
       (log(proposed.posterior.disps0) -
          log(current.posterior.disps0))^2 / (2*disp.proposal.scales0)) -
    disp.conditional.log.posterior(
      genes=genes, counts=counts, n=samples0,
      sample.means=sample.means0, means=current.posterior.means0,
      disps=current.posterior.disps0,
```

```
      prior.location=current.posterior.disp.prior.location,
      prior.scale=current.posterior.disp.prior.scale
    ) -
    (log(current.posterior.disps0) +
       (log(current.posterior.disps0) -
          log(proposed.posterior.disps0))^2 / (2*disp.proposal.scales0))

  current.posterior.disps0[replace] <- proposed.posterior.disps0[replace]
  accept.disps0[replace] <- accept.disps0[replace] + 1/chain.length

  # Metropolis updates for per-gene group 1 dispersions ####
  proposed.posterior.disps1 <- rlnorm(
    n=genes, meanlog=log(current.posterior.disps1), sdlog=sqrt.disp.proposal.scales1
  )

  replace <- log(runif(genes)) <=
    disp.conditional.log.posterior(
      genes=genes, counts=counts1, n=samples1,
      sample.means=sample.means1, means=current.posterior.means1,
      disps=proposed.posterior.disps1,
      prior.location=current.posterior.disp.prior.location,
      prior.scale=current.posterior.disp.prior.scale
    ) +
    (log(proposed.posterior.disps1) +
       (log(proposed.posterior.disps1) -
          log(current.posterior.disps1))^2 / (2*disp.proposal.scales1)) -
    disp.conditional.log.posterior(
      genes=genes, counts=counts1, n=samples1,
      sample.means=sample.means1,
      means=current.posterior.means1,
      disps=current.posterior.disps1,
      prior.location=current.posterior.disp.prior.location,
      prior.scale=current.posterior.disp.prior.scale
    ) -
    (log(current.posterior.disps1) +
       (log(current.posterior.disps1) -
          log(proposed.posterior.disps1))^2 / (2*disp.proposal.scales1))

  current.posterior.disps1[replace] <- proposed.posterior.disps1[replace]
  accept.disps1[replace] <- accept.disps1[replace] + 1/chain.length

  # Metropolis updates for per-gene group 2 dispersions ####
  proposed.posterior.disps2 <- rlnorm(
    n=genes, meanlog=log(current.posterior.disps2), sdlog=sqrt.disp.proposal.scales2
  )
  replace <- log(runif(genes)) <=
    disp.conditional.log.posterior(
      genes=genes, counts=counts2, n=samples2,
      sample.means=sample.means2, means=current.posterior.means2,
      disps=proposed.posterior.disps2,
      prior.location=current.posterior.disp.prior.location,
      prior.scale=current.posterior.disp.prior.scale
    ) +
```

```
    (log(proposed.posterior.disps2) +
       (log(proposed.posterior.disps2) -
          log(current.posterior.disps2))^2 / (2*disp.proposal.scales2)) -
    disp.conditional.log.posterior(
      genes=genes, counts=counts2, n=samples2,
      sample.means=sample.means2, means=current.posterior.means2,
      disps=current.posterior.disps2,
      prior.location=current.posterior.disp.prior.location,
      prior.scale=current.posterior.disp.prior.scale
    ) -
    (log(current.posterior.disps2) +
       (log(current.posterior.disps2) -
          log(proposed.posterior.disps2))^2 / (2*disp.proposal.scales2))

  current.posterior.disps2[replace] <-
    proposed.posterior.disps2[replace]
  accept.disps2[replace] <- accept.disps2[replace] + 1/chain.length

  # Gibbs update for prior location parameter for mean ####
  current.posterior.mean.prior.location <- rnorm(
    n=1,
    mean=prior.location.posterior.mean(
      hyperprior.mean=2.5, hyperprior.var=20,
      prior.scale=current.posterior.mean.prior.scale,
      parameters0=current.posterior.means0,
      parameters1=current.posterior.means1,
      parameters2=current.posterior.means2,
      z=current.posterior.indicators
    ),
    sd=prior.location.posterior.sd(
      hyperprior.var=20,
      prior.scale=current.posterior.mean.prior.scale,
      z=current.posterior.indicators
    )
  )

  # Metropolis update for prior scale parameter for mean ####
  proposed.posterior.mean.prior.scale <- rnorm(
    n=1, mean=current.posterior.mean.prior.scale, sd=mean.prior.scale.proposal.sd
  )

  if (log(runif(1)) <=
      mean.prior.scale.log.posterior(
        prior.scale=proposed.posterior.mean.prior.scale,
        prior.location=current.posterior.mean.prior.location,
        g=genes, means0=current.posterior.means0,
        means1=current.posterior.means1,
        means2=current.posterior.means2,
        z=current.posterior.indicators
      ) -
      mean.prior.scale.log.posterior(
        prior.scale=current.posterior.mean.prior.scale,
        prior.location=current.posterior.mean.prior.location,
```

```
      g=genes, means0=current.posterior.means0,
      means1=current.posterior.means1,
      means2=current.posterior.means2,
      z=current.posterior.indicators
    )) {
  current.posterior.mean.prior.scale <- proposed.posterior.mean.prior.scale
  accept.mean.prior.scale <- accept.mean.prior.scale + 1/chain.length
}

# Gibbs update for prior location parameter for dispersion ####
current.posterior.disp.prior.location <- rnorm(
  n=1,
  mean=prior.location.posterior.mean(
    hyperprior.mean=-2.5, hyperprior.var=2,
    prior.scale=current.posterior.disp.prior.scale,
    parameters0=current.posterior.disps0,
    parameters1=current.posterior.disps1,
    parameters2=current.posterior.disps2,
    z=current.posterior.indicators
  ),
  sd=prior.location.posterior.sd(
    hyperprior.var=2,
    prior.scale=current.posterior.disp.prior.scale,
    z=current.posterior.indicators
  )
)

# Metropolis update for prior scale parameter for dispersion ####
proposed.posterior.disp.prior.scale <- rnorm(
  n=1, mean=current.posterior.disp.prior.scale, sd=disp.prior.scale.proposal.sd
)

if (log(runif(1)) <=
    disp.prior.scale.log.posterior(
      prior.scale=proposed.posterior.disp.prior.scale,
      prior.location=current.posterior.disp.prior.location,
      g=genes, disps0=current.posterior.disps0,
      disps1=current.posterior.disps1,
      disps2=current.posterior.disps2,
      z=current.posterior.indicators
    ) -
    disp.prior.scale.log.posterior(
      prior.scale=current.posterior.disp.prior.scale,
      prior.location=current.posterior.disp.prior.location,
      g=genes, disps0=current.posterior.disps0,
      disps1=current.posterior.disps1,
      disps2=current.posterior.disps2,
      z=current.posterior.indicators
    )) {
  current.posterior.disp.prior.scale <- proposed.posterior.disp.prior.scale
  accept.disp.prior.scale <- accept.disp.prior.scale + 1/chain.length
}
```

```
    # Gibbs updates for per-gene mixture components ####
    current.posterior.indicators <- rbinom(
      n=genes, size=1,
      prob=posterior.indicator.probabilities(
        genes=genes, counts=counts, counts1=counts1, counts2=counts2,
        n=samples0, n1=samples1, n2=samples2, sample.means0=sample.means0,
        sample.means1=sample.means1, sample.means2=sample.means2,
        means0=current.posterior.means0, means1=current.posterior.means1,
        means2=current.posterior.means2, disps0=current.posterior.disps0,
        disps1=current.posterior.disps1, disps2=current.posterior.disps2,
        mean.prior.location=current.posterior.mean.prior.location,
        mean.prior.scale=current.posterior.mean.prior.scale,
        disp.prior.location=current.posterior.disp.prior.location,
        disp.prior.scale=current.posterior.disp.prior.scale,
        lambda=current.posterior.proportion
      )
    )

    # Gibbs update for mixture proportion ####
    current.posterior.proportion <- rbeta(
      n=1, shape1=1+sum(current.posterior.indicators),
      shape2=1+genes-sum(current.posterior.indicators)
    )

    # Update posterior samples ####
    if (iter/thin==round(iter/thin)) {
      posterior.means0[iter/thin,] <- current.posterior.means0
      posterior.means1[iter/thin,] <- current.posterior.means1
      posterior.means2[iter/thin,] <- current.posterior.means2
      posterior.disps0[iter/thin,] <- current.posterior.disps0
      posterior.disps1[iter/thin,] <- current.posterior.disps1
      posterior.disps2[iter/thin,] <- current.posterior.disps2
      posterior.mean.prior.location[iter/thin] <- current.posterior.mean.prior.location
      posterior.mean.prior.scale[iter/thin] <- current.posterior.mean.prior.scale
      posterior.disp.prior.location[iter/thin] <- current.posterior.disp.prior.location
      posterior.disp.prior.scale[iter/thin] <- current.posterior.disp.prior.scale
      posterior.indicators[iter/thin,] <- current.posterior.indicators
      posterior.proportion[iter/thin] <- current.posterior.proportion
    }

  }

  return(list(
    "chain.length"=chain.length, "thin"=thin, "inits"=inits,
    "mean.proposal.scales0"=mean.proposal.scales0,
    "mean.proposal.scales1"=mean.proposal.scales1,
    "mean.proposal.scales2"=mean.proposal.scales2,
    "disp.proposal.scales0"=disp.proposal.scales0,
    "disp.proposal.scales1"=disp.proposal.scales1,
    "disp.proposal.scales2"=disp.proposal.scales2,
    "mean.prior.scale.proposal.sd"=mean.prior.scale.proposal.sd,
    "disp.prior.scale.proposal.sd"=disp.prior.scale.proposal.sd,
    "accept.means0"=accept.means0,
```

```
    "accept.means1"=accept.means1,
    "accept.means2"=accept.means2,
    "accept.disps0"=accept.disps0,
    "accept.disps1"=accept.disps1,
    "accept.disps2"=accept.disps2,
    "accept.mean.prior.scale"=accept.mean.prior.scale,
    "accept.disp.prior.scale"=accept.disp.prior.scale,
    "posterior.means0"=posterior.means0,
    "posterior.means1"=posterior.means1,
    "posterior.means2"=posterior.means2,
    "posterior.disps0"=posterior.disps0,
    "posterior.disps1"=posterior.disps1,
    "posterior.disps2"=posterior.disps2,
    "posterior.mean.prior.location"=posterior.mean.prior.location,
    "posterior.mean.prior.scale"=posterior.mean.prior.scale,
    "posterior.disp.prior.location"=posterior.disp.prior.location,
    "posterior.disp.prior.scale"=posterior.disp.prior.scale,
    "posterior.indicators"=posterior.indicators,
    "posterior.proportion"=posterior.proportion
    ))

}
```

##### Log conditional posterior density functions

```
# Per-gene conditional log posterior mean function
# Takes vectors of per-gene sample means and mean and dispersion estimates
mean.conditional.log.posterior <- function(n, sample.means, means, disps, prior.location,
                                           prior.scale) {
  x <- 1/disps
  out <- rep(-1e20, length(means))
  index <- which(means>0)
  out[index] <-
    -n * (sample.means[index] + x[index]) * log(1 + means[index]*disps[index]) +
    (n*sample.means[index] - 1) * log(means[index]) -
    (log(means[index]) - prior.location)^2 / (2*prior.scale)
  return(out)
}


# Per-gene conditional log posterior dispersion function
# Takes vector of counts for each gene to calculate sum over log gamma for each gene, and
# vectors of per-gene sample means and mean and dispersion estimates
disp.conditional.log.posterior = function(genes, counts, n, sample.means, means, disps,
                                          prior.location, prior.scale) {
  x <- 1/disps
  out <- rep(-1e20,genes)
  index <- which(disps>0)
  lgammasum <- colSums(lgamma(counts + rep(x, each = nrow(counts))))
  out[index] <-
    lgammasum[index] -
    n * lgamma(x[index]) -
```

```r
    n * (sample.means[index] + x[index]) * log(1 + means[index]*disps[index]) +
    (n*sample.means[index] - 1) * log(disps[index]) -
    (log(disps[index]) - prior.location)^2 / (2*prior.scale)
  return(out)
}


# Function for sd of normal conditional posteriors for lognormal location parameters
# Takes vector of posterior indicators
prior.location.posterior.sd <- function(hyperprior.var, prior.scale, z) {
  return(sqrt(1 / (1/hyperprior.var + sum(1 + z) / prior.scale)))
}


# Function for mean of normal conditional posteriors for lognormal location parameters
# Takes vectors of mean or dispersion estimates and posterior indicators
prior.location.posterior.mean <- function(hyperprior.mean, hyperprior.var, prior.scale,
                                          parameters0, parameters1, parameters2, z) {
  return(
    (hyperprior.mean / hyperprior.var +
       sum((1-z)*log(parameters0) +
             z*(log(parameters1) + log(parameters2))) /
       prior.scale) / (1/hyperprior.var + sum(1+z)/prior.scale)
  )
}


# Conditional log posterior function for scale parameter of prior on mean
# Takes vectors of per-gene mean estimates and mixture components
mean.prior.scale.log.posterior <- function(prior.scale, prior.location, g, means0,
                                           means1, means2, z) {
  if (prior.scale <= 0) {
    return(-1e20)
  } else {
    return(
      (2 - (g + sum(z))/2) * log(prior.scale) -
        0.8 * prior.scale - sum(
          (1-z)*(log(means0) - prior.location)^2 +
            z*((log(means1) - prior.location)^2 +
                 (log(means2) - prior.location)^2)
        ) / (2*prior.scale)
    )
  }
}


# Conditional log posterior function for scale parameter of prior on dispersion
# Takes vectors of per-gene dispersion estimates and mixture components
disp.prior.scale.log.posterior <- function(prior.scale, prior.location, g, disps0,
                                           disps1, disps2, z) {
  if (prior.scale <= 0) {
    return(-1e20)
  } else {
```

```r
    return(
      (4 - (g + sum(z))/2) * log(prior.scale) -
        2 * prior.scale - sum(
          (1-z)*(log(disps0) - prior.location)^2 +
            z*((log(disps1) - prior.location)^2 +
                 (log(disps2) - prior.location)^2)
        ) / (2*prior.scale)
    )
  }
}


# Function for log posterior probability z=0
# Takes vector of counts for each gene to calculate sum over log gamma for each gene, and
# vectors of per-gene sample means and mean and dispersion estimates
pz0 <- function(genes, counts, n, sample.means, means, disps,
                mean.prior.location, mean.prior.scale,
                disp.prior.location, disp.prior.scale, lambda) {
  x <- 1/disps
  lgammasum <- colSums(lgamma(counts + rep(x, each = nrow(counts))))
  return(lgammasum -
           n * lgamma(x) -
           n * (sample.means + x) * log(1 + means * disps) +
           (n * sample.means - 1) * log(means * disps) +
           log(2*pi) +
           log(mean.prior.scale)/2 +
           log(disp.prior.scale)/2 -
           (log(means) - mean.prior.location)^2 /
             (2*mean.prior.scale) -
           (log(disps) - disp.prior.location)^2 /
             (2*disp.prior.scale) +
           log(1-lambda))
}


# Function for log posterior probability z=1
# Takes vector of counts for each gene to calculate sums over log gamma for each gene for
# each group, and vectors of per-gene sample means and mean and dispersion estimates for
# each group
pz1 = function(genes, counts1, counts2, n1, n2, sample.means1,
               sample.means2, means1, means2, disps1, disps2,
               mean.prior.location, mean.prior.scale,
               disp.prior.location, disp.prior.scale, lambda) {
  x1 <- 1/disps1
  x2 <- 1/disps2
  lgammasum1 <- colSums(lgamma(counts1 + rep(x1, each = nrow(counts1))))
  lgammasum2 <- colSums(lgamma(counts2 + rep(x2, each = nrow(counts2))))
  return(lgammasum1 -
           n1 * lgamma(x1) -
           n1 * (sample.means1 + x1) * log(1 + means1 * disps1) +
           (n1 * sample.means1 - 1) * log(means1 * disps1) +
           lgammasum2 -
           n2 * lgamma(x2) -
```

```r
          n2 * (sample.means2 + x2) * log(1 + means2 * disps2) +
          (n2 * sample.means2 - 1) * log(means2 * disps2) -
          ((log(means1) - mean.prior.location)^2 +
             (log(means2) - mean.prior.location)^2) / (2*mean.prior.scale) -
          ((log(disps1) - disp.prior.location)^2 +
             (log(disps2) - disp.prior.location)^2) / (2*disp.prior.scale) +
          log(lambda))
}


# Function to calculate probabilities for posterior Bernoulli distributions for mixture
# components (exponentiates and normalises calculated probabilities)
posterior.indicator.probabilities <- function(genes, counts0, counts1, counts2,
                                              n, n1, n2, sample.means0, sample.means1,
                                              sample.means2, means0, means1, means2,
                                              disps0, disps1, disps2,
                                              mean.prior.location, mean.prior.scale,
                                              disp.prior.location, disp.prior.scale,
                                              lambda) {
  return(1 / (1 + exp(pz0(genes=genes,
                          counts=counts0,
                          n=n,
                          sample.means=sample.means0,
                          means=means0,
                          disps=disps0,
                          mean.prior.location=mean.prior.location,
                          mean.prior.scale=mean.prior.scale,
                          disp.prior.location=disp.prior.location,
                          disp.prior.scale=disp.prior.scale,
                          lambda=lambda) -
                        pz1(genes=genes,
                            counts1=counts1,
                            counts2=counts2,
                            n1=n1,
                            n2=n2,
                            sample.means1=sample.means1,
                            sample.means2=sample.means2,
                            means1=means1,
                            means2=means2,
                            disps1=disps1,
                            disps2=disps2,
                            mean.prior.location=mean.prior.location,
                            mean.prior.scale=mean.prior.scale,
                            disp.prior.location=disp.prior.location,
                            disp.prior.scale=disp.prior.scale,
                            lambda=lambda)
  )
  )
  )
}
```

##### Functions to compute posterior tail probabilities that log fold changes in mean or dispersion are less than or equal to a given minimum value

```
# Highest density interval function
# Uses code from hdi() from HDInterval package, but removed sorting part so can sort data
# once in tail probability function instead of sorting for every iteration
hdi <- function(x, credMass) {
  n <- length(x)
  exclude <- n - floor(n * credMass)
  low.poss <- x[1:exclude]
  upp.poss <- x[(n - exclude + 1):n]
  best <- which.min(upp.poss - low.poss)
  result <- c(low.poss[best], upp.poss[best])
  return(result)
}

# Function to calculate tail probability given min log fold change m
hpd.pval <- function(x, m=0) {
  x <- sort.int(x, method='quick')

  y1 <- numeric(9)
  for (t in 1:9) {
    y1[t] <- sign(hdi(x, credMass=1 - t * 0.1)[1] - m) ==
      sign(hdi(x, credMass=1 - t * 0.1)[2] + m)
    if (y1[t] == 1) {break}
  }
  if (sum(y1) == 0) {x1 <- 9} else {x1 <- min(which(y1 == 1)) - 1}
  iter <- x1 * 0.1

  y2 <- numeric(9)
  for (t in 1:9) {
    y2[t] <- sign(hdi(x, credMass=1 - (iter + t * 0.01))[1] - m) ==
      sign(hdi(x, credMass=1 - (iter + t * 0.01))[2] + m)
    if (y2[t] == 1) {break}
  }
  if (sum(y2) == 0) {x2 <- 9} else {x2 <- min(which(y2 == 1)) - 1}
  iter <- iter + x2 * 0.01

  y3 <- numeric(9)
  for (t in 1:9) {
    y3[t] <- sign(hdi(x, credMass=1 - (iter + t * 1e-3))[1] - m) ==
      sign(hdi(x, credMass=1 - (iter + t * 1e-3))[2] + m)
    if (y3[t] == 1) {break}
  }
  if (sum(y3) == 0) {x3 <- 9} else {x3 <- min(which(y3 == 1)) - 1}
  iter <- iter + x3 * 1e-3

  y4 <- numeric(9)
  for (t in 1:9) {
    y4[t] <- sign(hdi(x, credMass=1 - (iter + t * 1e-4))[1] - m) ==
      sign(hdi(x, credMass=1 - (iter + t*1e-4))[2] + m)
    if (y4[t] == 1) {break}
  }
```

```r
  if (length(x) <= 1e4) {
    if (sum(y4) == 0) {x4 <- 10} else {x4 <- min(which(y4 == 1))}
    iter <- iter + x4 * 1e-4
  }
  else {
    if (sum(y4) == 0) {x4 <- 9} else {x4 <- min(which(y4 == 1)) - 1}
    iter <- iter + x4 * 1e-4

    y5 <- numeric(9)
    for (t in 1:9) {
      y5[t] <- sign(hdi(x, credMass=1 - (iter + t * 1e-5))[1] - m) ==
        sign(hdi(x, credMass=1 - (iter + t * 1e-5))[2] + m)
      if (y5[t] == 1) {break}
    }
    if (sum(y5) == 0) {x5 <- 10} else {x5 <- min(which(y5 == 1))}
    iter <- iter + x5 * 1e-5
  }

  return(iter)
}
```

#### Bayesian FDR computation

```r
# Based on BFDR() from ShrinkBayes package
bfdr <- function(x) {
  ord <- sort(x, decreasing=T, index.return=T)
  bfdr <- cumsum(1-ord$x)/(1:length(x))
  bfdr[ord$ix] <- bfdr
  return(bfdr)
}
```
