## Additional file 4: Methods comparisons for "Identification of differentially distributed gene expression and distinct sets of cancer-related genes identified by changes in mean and variability"

### Additional file 3 for *Identification of differentially distributed gene expression and distinct sets of cancer-related genes identified by changes in mean and variability*: R code for differential expression, dispersion and distribution methods comparisons

#### Normalisation

```
libsizes <- colSums(counts)
nf <- calcNormFactors(counts, method="TMM") # normalisation factors for edgeR, limma
els <- nf * libsizes # effective library sizes for baySeq
sf <- els / exp(mean(log(libsizes))) # size factors for DESeq2, MDSeq
norm.counts <- t(t(counts) / sf) # normalised count matrix used for diffVar, HM
```

#### edgeR

```
design <- model.matrix(~group) # group is a factor of group labels
dat.edgeR <- DGEList(counts=counts, norm.factors=nf, group=group)
dat.edgeR <- estimateDisp(dat.edgeR, design)
fit.edgeR <- glmQLFit(dat.edgeR, design)
test.edgeR <- glmQLFTest(fit.edgeR)
```

#### DESeq2

```
dat.DESeq2 <- DESeqDataSetFromMatrix(countData=counts, colData=data.frame(group),
                                     design=~group)
sizeFactors(dat.DESeq2) <- sf
fit.DESeq2 <- DESeq(dat.DESeq2, minReplicatesForReplace=Inf)
res.DESeq2 <- results(fit.DESeq2, cooksCutoff=F, alpha=0.05)
```

#### limma-voom

```
dat.voom <- voom(dat.edgeR) # uses same data object as edgeR
fit.voom <- lmFit(dat.voom, design)
res.voom <- eBayes(fit.voom)
```

#### baySeq

```
dat.baySeq <- new('countData', data=counts, replicates=group,
                  groups=list(NDE=rep(1,length(group)), DE=as.numeric(group)))
dat.baySeq@annotation <- data.frame(name=1:nrow(dat.baySeq@data))
libsizes(dat.baySeq) <- els
dat.baySeq <- getPriors.NB(dat.baySeq, samplesize=10000)
dat.baySeq <- getLikelihoods(dat.baySeq)
res.baySeq <- topCounts(dat.baySeq, group='DE', number=Inf)[
  order(topCounts(dat.baySeq, group='DE', number=Inf)$name), ]
```

#### MDSeq

```
contrasts <- get.model.matrix(group)
fit.MDSeq.zi <- MDSeq(counts, offsets=sf, contrast=contrasts)
res.MDSeq.zi <- extract.ZIMD(fit.MDSeq.zi, compare=list(A="1",B="2"))
# performs both differential expression and differential dispersion tests
```

#### GAMLSS

```
# Code modified from https://github.com/Vityay/ExpVarQuant/blob/master/ExpVarQuant.R

sink(file = "diversionOfOutput.txt", append = TRUE)
dat.edgeR$CPM <- cpm.DGEList(dat.edgeR) # uses same data object as edgeR
ofs <- log(dat.edgeR$samples$lib.size * dat.edgeR$samples$norm.factors)
dat.edgeR$samples$offset <- ofs
gene_i <- seq_along(dat.edgeR$counts[,1])

gamlss_NB <- lapply(gene_i, function(i) {
  dat <- data.frame(x = dat.edgeR$samples$group,
                    y = dat.edgeR$counts[i,],
                    ofs = dat.edgeR$samples$offset)
  dat$x <- relevel(dat$x, ref = c("1"))
  m0 <- tryCatch(gamlss(fo = y ~ 0+x+offset(ofs), sigma.fo = ~ 0+x, data=dat,
                        family = NBI(), sigma.start = 0.1, n.cyc = 100),
                 warning= function(w) NULL, error= function(e) NULL)
  m1 <- tryCatch(gamlss(fo = y ~ 0+x+offset(ofs), sigma.fo = ~ 1, data=dat,
                        family = NBI(), sigma.start = 0.1, n.cyc = 100),
                 warning= function(w) NULL, error= function(e) NULL)
  res <- data.frame(CV.1 = NA,
                    CV.2 = NA,
                    LR.cv = NA,
                    p.cv = NA)
  if(!any(sapply(list(m0,m1), is.null)))
  {
    res$CV.1  = sqrt(exp(m0$sigma.coefficients)[[1]])
    res$CV.2 = sqrt(exp(m0$sigma.coefficients)[[2]])
    res$LR.cv = log2(sqrt(exp(m0$sigma.coefficients)[[2]]) /
                       sqrt(exp(m0$sigma.coefficients)[[1]]))
    res$p.cv = pchisq(2*(logLik(m0)-logLik(m1)), df=m0$df.fit-m1$df.fit, lower=F)[[1]]
  }
  res
})
closeAllConnections()

gamlss_NB <- do.call(rbind, gamlss_NB)
rownames(gamlss_NB) <- rownames(dat.edgeR$counts)[gene_i]
res_clean <- na.omit(gamlss_NB)
```

#### diffVar

```
fit.diffVar <- varFit(norm.counts, design=design, coef=c(1,2))
res.diffVar <- topVar(fit.diffVar, coef=2, number=nrow(norm.counts), sort=F)
```

#### Hierarchical model

```
HM <- ln_hmm_adapt_3_chains(counts=t(norm.counts), groups=group)

# Differential expression:
mean.diff.HM <- unname(log(as.matrix(HM$means1)) - log(as.matrix(HM$means2)))
# posterior samples of between-group differences in means
p.mean.HM <- apply(mean.diff.HM, 2, hpd.pval)
# tail probabilities of no differences in means

# Differential dispersion:
disp.diff.HM <- unname(log(as.matrix(HM$disps1)) - log(as.matrix(HM$disps2)))
# posterior samples of between-group differences in dispersions
p.disp.HM <- apply(disp.diff.HM, 2, hpd.pval)
# tail probabilities of no differences in dispersions

# Differential distribution:
prob.HMM <- unname(colMeans(as.matrix(HM$indicators)))
# raw posterior probabilities of differential distribution
bfdr.HMM <- bfdr(prob.HMM)
# Bayesian FDR method for setting posterior probability threshold
post.prop.HMM <- mean(as.matrix(HM$proportion))
# estimated proportion of differentially distributed genes
thr.HMM <- sort(prob.HMM, decreasing=T)[round(nrow(counts) * post.prop.HMM)]
# posterior probability threshold to give appropriate proportion of differentially
# distributed genes for alternative posterior threshold method
```
